# TIDE: Tractography-Informed Dose Estimation for individualised TMS intensity

**DOI:** 10.64898/2026.08.20.746040

**Authors:** Marco Tagliaferri, Luigi Cattaneo, Carlo Miniussi, Arianna Brancaccio

**Author notes:** Corresponding author. Marco Tagliaferri, PhD Candidate in Cognitive and Brain Sciences, Department of Psychology and Neuroscience - DIPseN, University of Trento, E-mail (permanent).

## Abstract

Transcranial magnetic stimulation (TMS) is commonly dosed by setting stimulation intensity as a fixed percentage of the resting motor threshold (RMT), although a motor-derived intensity may not produce comparable neural recruitment across non-motor targets. We present TIDE (Tractography-Informed Dose Estimation), an open-source, SimNIBS-based pipeline designed to derive individualised stimulation intensities for non-motor white-matter targets. TIDE combines individual RMT measurements, finite-element electric-field modelling and diffusion MRI tractography to rescale the stimulation intensity according to the geometry and stimulation efficiency of the pathway of interest. Specifically, it computes the activating function along subject-specific streamlines and estimates the stimulator output, expressed as a percentage of maximum stimulator output, required for the target pathway to reach the activation level produced in the corticospinal tract at RMT. In an independent dataset of 19 participants, in which stimulation had been dosed conventionally as a fixed percentage of RMT, the relative difference between delivered and TIDE-estimated intensity was associated with the magnitude of TMS-induced behavioural effects at two frontal aslant tract (FAT) stimulation sites, while the delivered intensity alone was not. TIDE therefore extends conventional E-field dosing from cortical field magnitude to subject-specific pathway geometry, providing a method to move beyond the assumption of homogeneous pathway engagement while accounting for inter-individual variability in pathway-specific stimulation efficiency.

## 1. Introduction

The response elicited by transcranial magnetic stimulation (TMS) is determined by the interaction between the induced electric field (E-field) and the subject-specific anatomy of the stimulated brain. Stimulation intensity, coil geometry, position and orientation, cortical folding and tissue geometry shape the magnitude, direction and spatial distribution of the induced E-field (Dannhauer et al., 2024; Lee et al., 2018; Thielscher et al., 2011). Neural recruitment then depends on how this field interacts with the architecture of the underlying neural elements (Aberra et al., 2020; Rattay, 1986, 1989; Roth & Basser, 1990). Recruitment therefore unfolds in two stages: the stimulation parameters and individual anatomy set the induced E-field, which in turn interacts with the geometry of the underlying axonal structures to determine neural recruitment. Because neural recruitment depends on the interaction between the induced field and the underlying neural architecture, even comparable induced E-fields across participants, or across targets within the same participant, may result in different levels of pathway engagement, defined here as the recruitment of the targeted neural pathway.

To account for individual and regional variability in pathway engagement, individualised dosing scales stimulation intensity to each participant’s target anatomy rather than relying on standard motor-threshold (MT) transfer. Conceptually, the stimulation dose is defined as the induced E-field magnitude at the target required to achieve equivalent neural engagement across individuals. In practice, this dose is implemented by adjusting the maximun stimulator output (%MSO) for each participant to produce the desired E-field at the target.

The importance of controlling how the targeted neural system is engaged in each individual is particularly evident in TMS combined with electroencephalography (EEG). TMS-evoked potentials (TEPs) show substantial between-subject variability in amplitude, latency and spatial distribution, even when the same cortical region is targeted (ter Braack et al., 2018). Part of this variability reflects fluctuations in the functional state of the stimulated local neural system (Desideri et al., 2019; Zrenner et al., 2018), but TEPs are also shaped by the physical parameters of stimulation including coil orientation and induced-current direction (Granö et al., 2025; Guidali et al., 2023; Mancuso et al., 2024; Parmigiani et al., 2025). The measured response therefore depends on how the target is engaged rather than on the stimulated region alone. This illustrates a general problem that extends beyond TMS-EEG: the effects produced by TMS, whether measured as electrophysiological responses or as behavioural changes, cannot be interpreted independently of the effective dose delivered to the targeted neural structures.

Stimulation intensity is a major component of this uncertainty. Outside the primary motor cortex (M1), TMS dose is commonly prescribed as a fixed percentage of the resting MT (RMT), e.g., 120% of RMT. This practice transfers an intensity calibrated in the motor system to anatomically distinct cortical targets and implicitly assumes that the relationship between motor-referenced stimulator output and neural recruitment is sufficiently comparable across cortical regions. Measuring the RMT in every participant individualises the starting point of this transfer, but not the transfer itself, which depends on anatomical features that vary independently of the MT. However, a given stimulator output does not produce a uniform E-field throughout the brain. For a given coil geometry field magnitude, direction and spatial distribution depend on scalp-to-cortex distance, cortical folding, tissue geometry, coil position and orientation (Dannhauer et al., 2024; Lee et al., 2018; Thielscher et al., 2011). Early approaches partly addressed this by adjusting the RMT for scalp-to-cortex distance (Stokes et al., 2005, 2007), an approximation now largely superseded by subject-specific E-field modelling. In this regard, simulations comparing motor and prefrontal stimulation indicate that intensities derived from M1 do not necessarily produce comparable E-fields or cellular responses in the dorsolateral prefrontal cortex (Turi et al., 2022).

This limitation does not remove the value of RMT as a calibration measure. Motor evoked potentials are variable because the recorded response is shaped by fluctuations across cortical, corticospinal and spinal components of the motor system (Ammann et al., 2020; Kiers et al., 1993; Rösler et al., 2008; Spampinato et al., 2023). Nevertheless, they provide a directly observable, subject-specific physiological response to a TMS pulse. An equivalent response is not readily available for non-motor cortical targets. RMT therefore remains a useful empirical anchor. Its validity as a calibration measure, however, does not imply that the corresponding stimulator output can be transferred to another region as though the same percentage of RMT represented the same effective neural dose.

To address this problem, previous works have developed subject-specific E-field modelling. E-field-based dosing methods commonly estimate the E-field produced at M1 at an intensity derived from the individual MT and adjust the stimulator output to obtain the same E-field magnitude at a non-motor target (Caulfield et al., 2021; Dannhauer et al., 2024; Numssen et al., 2024). Compared with fixed RMT scaling, this approach accounts for individual differences in anatomy and for regional variation in the relationship between stimulator output and cortical field strength. E-field-based dosing addresses this residual at the level of cortical field magnitude, and therefore provides a more comparable physical measure of stimulation across targets and participants.

However, matching the cortical E-field magnitude at the target to that induced at M1 does not guarantee recruitment of the intended neuronal population or an equivalent level of neural activation. E-field-based dosing remains sensitive to the definition and localisation of the target, the accuracy of the underlying simulations, and the assumption that stimulation thresholds are comparable across cortical regions and stimulation protocols (Numssen et al., 2024). A further limitation is that cortical field magnitude alone does not describe how the field interacts with the geometry of the underlying axons. Neural polarisation depends on the orientation of the induced field relative to neuronal processes and on spatial changes in the field component parallel to the axonal trajectory (Aberra et al., 2020; Rattay, 1986, 1989; Roth & Basser, 1990). Two cortical targets exposed to a similar E-field magnitude may therefore differ in their capacity to recruit the white-matter pathways that terminate in or pass beneath them. Accounting for the geometry of these pathways therefore provides a more biologically relevant basis for translating a motor-derived calibration to non-motor targets.

In this light, we developed Tractography-Informed Dose Estimation (TIDE), an open-source pipeline that combines individual RMT measurements, finite-element E-field simulations and diffusion MRI tractography. TIDE uses the corticospinal tract beneath the M1 hotspot as the calibration pathway and the subject’s RMT as the empirical dose anchor. For the target pathway, TIDE evaluates multiple candidate coil positions. Each position is defined by where the centre of the coil is placed on the scalp and by how the coil is oriented in three dimensions, including handle rotation and tilt relative to the local scalp surface. For each candidate position, TIDE simulates the E-field and evaluates it at successive points along the streamlines of both the corticospinal tract and the tractography-defined target bundle. TIDE then computes the activating function, defined as the spatial derivative of the E-field component parallel to each streamline, and identifies both the stimulator output and coil position required for the target pathway to reproduce the bundle-level activating function generated in the corticospinal tract at RMT. TIDE thus returns an individualised stimulation configuration, comprising the target-specific intensity and coil position.

TIDE thus extends conventional E-field dosing beyond cortical field magnitude to the geometry of the pathway intended to be engaged. This approach rests on the central assumption that the effect of TMS on these pathways is mediated by direct activation of their white-matter axons, the mechanism that in the corticospinal tract underlies the D-wave, rather than by indirect trans-synaptic recruitment. At suprathreshold intensities such as the 120% of RMT used in the present validation, the induced field directly activates axons at the juxtacortical white matter and grey-white matter junction (Aberra et al., 2020; Di Lazzaro & Rothwell, 2014; Siebner et al., 2022). To assess the biological relevance of TIDE estimates, we retrospectively applied the pipeline to an independent dataset in which single-pulse TMS delivered to tractography-defined endpoints of the left FAT produced site-specific behavioural effects (Tagliaferri et al., 2023). We tested whether participants whose delivered intensities was closer to their TIDE-estimate intensity showed larger sham-corrected behavioural shifts. We further tested whether tract-specific stimulation intensities could be generalised across individuals, by estimating the between-subject variability of the TIDE-recommended intensity across 13 white-matter targets.

## 2. Methods

TIDE is implemented as a thin layer over SimNIBS 4.5 (Thielscher et al., 2015): the head-model pipeline, the FEM solver, and the coil-optimisation routine are inherited from SimNIBS without modification, and the contribution of TIDE lies in the streamline-based activating-function analysis and the calibration inversion that translate the SimNIBS-computed E-field into a target-specific stimulator output.

The TIDE pipeline accepts three subject-specific inputs (a T1-weighted MRI volume, one tractogram per bundle of interest, and the subject’s RMT measured on the abductor pollicis brevis or first dorsal interosseous, expressed as a %MSO) and returns the estimated stimulator intensity required to produce the same biological activation threshold at any other target bundle. The estimator relies on no population priors, no anisotropic conductivity model, and no calibration beyond the RMT itself.

### 2.1 Pipeline overview

TIDE provides two primary operational workflows based on the same numerical core. The estimation workflow computes the recommended intensity for a single calibration–target pair, I_TIDE, as defined in Eq. (5). The grid-search workflow repeats the same estimation across candidate cortical positions associated with the target tractogram and returns a cortical map of I_TIDE values. Implementation details and output artefacts are described in Sections 2.10.1–2.10.3.

### 2.2 Notations and conventions

Throughout this work, coordinates are expressed in RASMM space. The T1-weighted volume is the anatomical reference for tractogram loading via DIPY (Garyfallidis, Brett, Amirbekian, Rokem, van der Walt, et al., 2014) into a ‘StatefulTractogram’; SimNIBS meshes are kept in the same space, which removes the need for inter-space resampling of streamline coordinates at field-sampling time.

Three-unit conventions are fixed for the remainder of the manuscript. The stimulator current slew rate, dI/dt, is expressed in A/s; the device maximum dI/dt is a coil-specific constant supplied by the manufacturer (1.61×10⁸ A/s for the MagVenture C-B60 model used in our reference subject). All finite-element simulations are run at a reference dI/dt of UNIT_DIDT = 1×10⁶ A/s so that any reported activating function value can be read as an intrinsic efficiency in V/m² per A/µs, and rescaled to any clinical intensity by a single multiplication. The subject’s RMT, the delivered intensity I_delivered, and the estimated target intensity I_TIDE are all expressed as percentages of the device maximum dI/dt (“% max output”), which places the measured and the estimated quantities on the same axis.

The signed activating function is preserved upstream, and the consumers that operate on activation magnitude (the contiguous-threshold estimator of Section 2.6 and the cross-streamline aggregator of Section 2.7) apply |AF| explicitly at the point of use rather than discarding polarity earlier. The storage conventions for the signed and magnitude forms across the streamline and volumetric outputs are given in Supplementary Methods S4.

We adopt the following shorthand. **E**(*s*) denotes the E-field vector at arc-length *s* along a streamline; *T* denotes the unit tangent taken from the Frenet-Serret frame; AF(*s*) denotes the activating function defined in Eq. (1) below; AFCST and AFtarget denote the cross-streamline aggregates of Section 2.7 evaluated on the calibration and target bundles; and ROI denotes the spherical region of analysis centred on the user-supplied cortical coordinates with a configurable radius (30 mm by default).

Stimulator intensity is denoted by I and expressed in % of maximum stimulator output (% max output), the standard TMS unit. Two intensities are distinguished. I_delivered is the intensity applied in the lab (in the present FAT validation cohort, RMT × 120 %); I_TIDE is the intensity recommended by the calibration inversion of Eq. (5), reported in raw and clamped form (*I*_*TIDE*, clamped; Supplementary Methods S9). The signed per-subject mismatch is *δI* = *I*delivered − *I*TIDE; the unsigned relative mismatch is |Δ*I*|/*I*TIDE. The Stimulation Efficiency Index SEI = AF_target / AF_CST = RMT / I_TIDE is dimensionless.

### 2.3 Tractogram requirements and recommended preprocessing

The TIDE pipeline operates on streamlines that are taken as faithful samples of the underlying fibre geometry. The accuracy of the recovered activating function is bounded by the accuracy of those streamlines, so the recommendations below describe a reconstruction protocol that we have validated against the assumptions of Sections 2.4–2.6.

Automated reconstruction retains streamlines that are anatomically implausible, and TIDE evaluates the activating function on whatever geometry the tractogram supplies. Every bundle in the multi-bundle analysis of Section 3.2 was therefore inspected against the anatomical reference and manually refined before analysis, and the cortical coordinates used to seed coil optimisation were taken from the refined bundles. Both steps were performed in TractEdit (v3.4.7; Tagliaferri & Cattaneo, 2026).

The full acquisition, preprocessing, fibre-orientation, reconstruction, and bundle-coverage protocol is given in Supplementary Methods S1.

### 2.4 Activating Function on white-matter streamlines

The cable-equation derivation of the activating function for an unmyelinated fibre embedded in an external E-field is due to Rattay (Rattay, 1986, 1989). For a streamline parameterised by arc length s, with unit tangent *T*(*s*) taken from the Frenet–Serret frame and induced field *E*(*r*(*s*)) sampled at position *r*(*s*) (**Figure 2**):

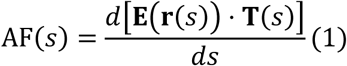

**Figure 1.**
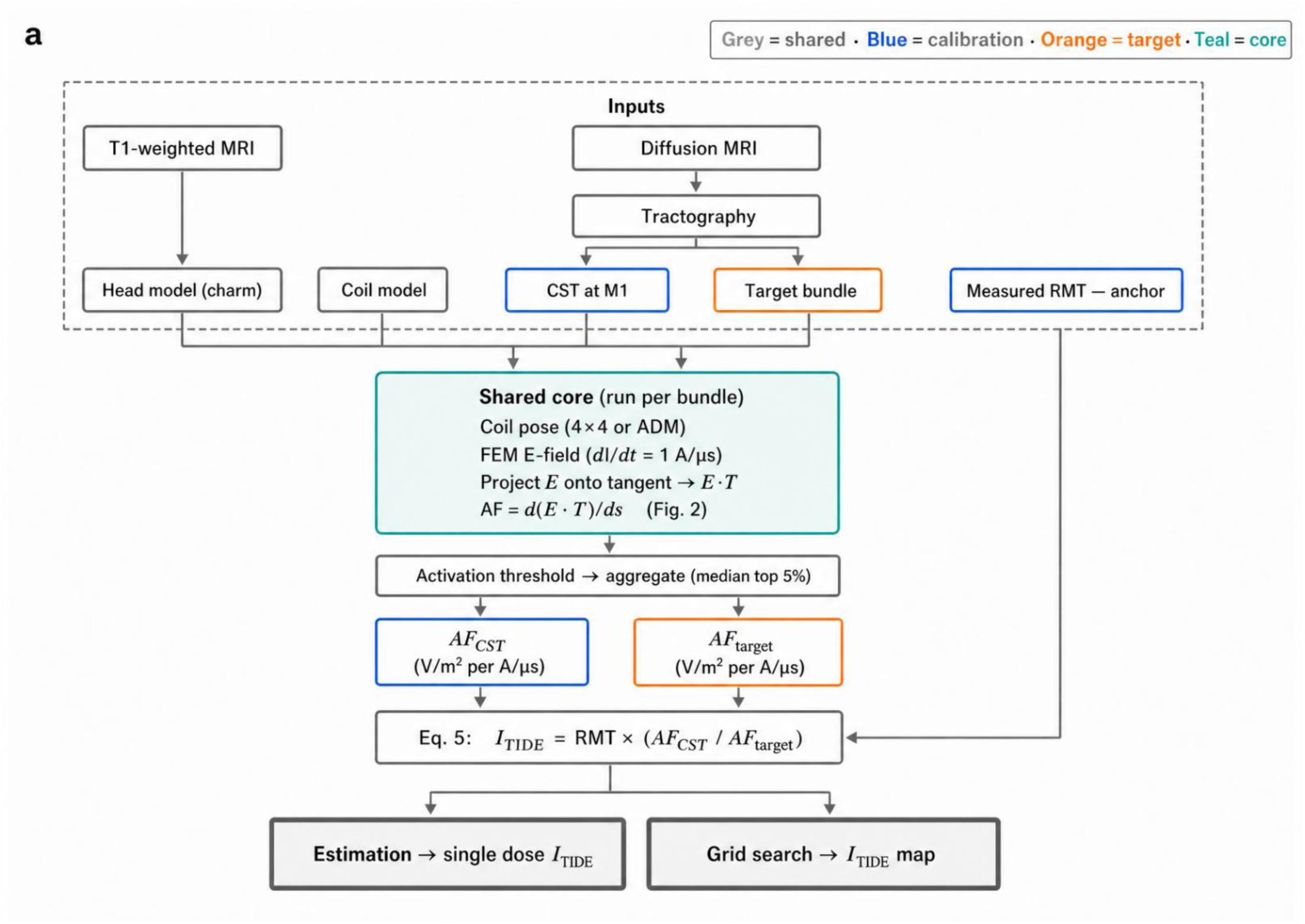
The TIDE pipeline workflow. Shared inputs (grey) are a T1-weighted MRI, converted to a SimNIBS head model with charm, a coil model, and diffusion MRI, from which tractography reconstructs two bundles: the corticospinal tract sampled at the motor hotspot (CST at M1; calibration, blue) and the target bundle (orange). A shared core (teal) is run once per bundle: it places the coil pose (a supplied 4×4 matrix or an ADM optimisation), solves the FEM E-field at unit rate of change of current (dI/dt = 1 A/µs), projects the field onto each streamline tangent (E·T), and computes the activating function as its arc-length gradient, AF = d(E·T)/ds (Fig. 2). Per-streamline activation thresholds are aggregated as the median of the top 5 % to give one efficiency value per bundle. The target dose follows from Eq. 5, ITIDE = RMT × (AFCST / AFtarget), where the measured resting motor threshold (RMT) is the only empirical anchor. The same core supports two outputs: a single-target estimation returning one dose ITIDE, and a grid search returning a spatial map of ITIDE over candidate scalp positions.

**Figure 2.**
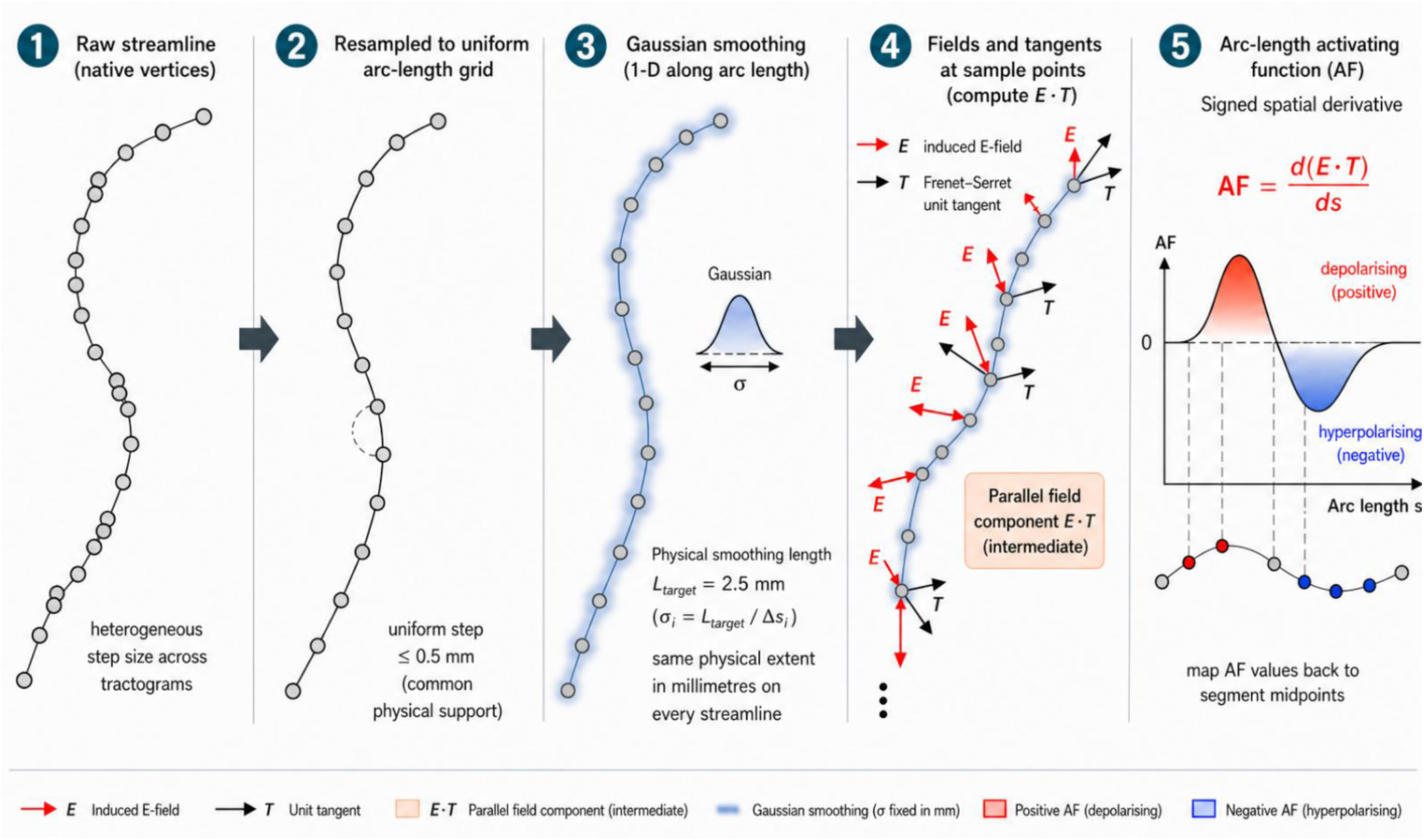
Arc-length formulation of the Activating Function (AF) in TIDE. Per-streamline computation of the gradient Activating Function *AF* = *d*(*E* · *T*)/*ds* (*Eq*. 1) **(1)** A raw streamline is stored on its native vertices, whose step size is heterogeneous across tractograms and reconstruction algorithms. **(2)** The streamline geometry and its co-located E-field vectors are jointly resampled by linear interpolation onto a uniform arc-length grid, using the smallest number of equal intervals that brings the step to ≤ 0.5 mm (common physical support). **(3)** The resampled coordinates and field are smoothed with a one-dimensional Gaussian whose extent is fixed in millimetres rather than in points: the support-relative standard deviation is *σi* = *Ltarget*/*Δsi* (Eq. 2), with physical smoothing length L_target = 2.5 mm by default, so the low-pass filter has the same spatial extent on every streamline regardless of native resolution. **(4)** On the smoothed curve, the Frenet–Serret unit tangent T (black) is computed and the induced E-field E (red) is projected onto it to form the parallel component E·T, an intermediate quantity. **(5)** The signed derivative of E·T with respect to arc length is taken as a forward difference between adjacent samples, assigned to the interval midpoint and divided by the smoothed interval length (second-order accurate), then mapped back to the midpoint of each original streamline segment. Positive AF (red) denotes depolarising drive, negative AF (blue) hyperpolarising; the AF is stored signed, and magnitude consumers apply |AF| at the point of use.

This term is the spatial gradient of the parallel field component along the fibre. It carries the contribution that dominates where the projection of E onto the fibre direction changes rapidly with distance, for example near the grey-white interface and at sharp turns in the white-matter geometry. Because the unit tangent is recomputed as the streamline curves, the derivative in Eq. (1) already carries the effect of local curvature on the alignment between field and fibre, and no separate curvature term is added.

The numerical implementation departs from the standard formulation in one respect that is, to our knowledge, not addressed in earlier AF-based TMS modelling work. Tractograms produced by different reconstruction algorithms (deterministic streamline tracking, probabilistic tracking, surface-seeded variants) deliver streamlines with heterogeneous mean step sizes, both between subjects and within a single bundle. A Gaussian filter applied with the same σ to every streamline therefore enforces the same kernel in points, not in millimetres. The physical smoothing length becomes a function of the local step size, which under-smooths streamlines with coarse steps and over-smooths streamlines with fine steps. The along-fibre gradient in Eq. (1) is sensitive to this inhomogeneity, because it is a numerical derivative of the field sampled along the streamline coordinates and is therefore amplified by any non-uniformity in the spatial low-pass response.

We remove this dependence on the input sampling by evaluating the activating function on a common physical support instead of on the native streamline vertices (Figure 2). For each streamline, the geometry and the co-located field vectors are jointly resampled by linear interpolation onto a uniform arc-length grid derived from the cumulative Euclidean length of the streamline, using the smallest number of equal intervals that brings the step to 0.5 mm or below, subject to a floor of three intervals for very short streamlines. The realised step *Δs*_*i* is therefore uniform within a streamline and at most 0.5 mm, which is why Eq. (2) is written per streamline. The resampled coordinates and the resampled field are then smoothed with a one-dimensional Gaussian whose extent is fixed in millimetres, under a nearest-value boundary policy, using the support-relative standard deviation,

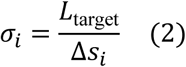

where *L*_*target* is the physical smoothing length (2.5 mm by default) and *Δs*_*i* is the realised resampling step of streamline ‘i’, so the low-pass filter has the same spatial extent in millimetres on every streamline irrespective of the native reconstruction resolution. The Frenet–Serret frame is computed on the smoothed, resampled coordinates with DIPY’s ‘frenet_serret’ routine (Garyfallidis, Brett, Amirbekian, Rokem, Van Der Walt, et al., 2014); the parallel component *E* · *T* is formed on this support; and its signed derivative with respect to arc length is taken as the forward difference between adjacent grid samples, assigned to the midpoint of the interval that separates them and divided by the smoothed interval length wherever that length is numerically non-zero; the resulting operator is second-order accurate at the point to which it is assigned.

Streamlines with fewer than four distinct arc-length positions are dropped: the gradient stencil needs at least two consecutive intervals to evaluate *dE* · *T*/*ds*, the arc-length resampling and Gaussian smoothing need a minimal support, and tractograms commonly contain short stubs that violate these requirements. The test is applied after duplicate positions are collapsed, so a streamline carrying repeated coincident vertices is treated according to the geometry it actually describes rather than the number of rows it stores. We treat the four-point cutoff as a single, physically motivated quality criterion.

The output of the AF computation is a triple of parallel arrays per surviving streamline: midpoint coordinates, signed AF values, and native segment lengths in millimetres. Each AF value is carried at the midpoint of the original streamline segment onto which it is mapped back from the uniform arc-length support, so the scalar sits between the two vertices that bound its segment. These midpoints, and not the original vertices, are the support on which the contiguous-activation criterion of Section 2.6 is subsequently evaluated; the criterion accumulates lengths in millimetres rather than point counts, so that the activation-length threshold remains a physical quantity and is portable across tractograms of different resolutions. The native segment lengths are carried alongside the AF values into the TRK outputs, where they support offline re-analysis at the original streamline resolution.

A small number of streamlines may fail the Frenet-Serret computation at runtime (numerical degeneracy on near-colinear segments, for example) or arrive with a mismatch between the number of streamline points and the number of sampled field vectors. The surviving population is therefore the largest set of streamlines for which Eq. (1) and Eq. (2) are well-defined; in practice, on the reference subject, fewer than 1% of streamlines are dropped at this stage. An optional angular-deviation filter is available upstream of the AF computation, controlled by options.max_angular_deviation_deg. When a positive threshold is set, a streamline is dropped in full if the angle between any two consecutive tangent vectors inside the ROI exceeds it. This targets streamlines that curl back on themselves at cortical endpoints, where the tangent, and therefore the gradient term of Eq. (1), is unreliable; the whole streamline is removed rather than the offending segment, because a corrupted tangent invalidates the gradient stencil along the affected span. The filter is applied after field sampling and before Eq. (1). Its default is 0.0, which disables it, and both the behavioural validation of Section 2.12 and the multi-bundle reference table were produced with the filter disabled.

### 2.5 Electrical field simulation and sampling

The T1-weighted volume is segmented into skin, skull, cerebrospinal fluid, grey- and white-matter compartments with charm, from which the tetrahedral head mesh is generated. The induced E-field is simulated with SimNIBS 4.5 (Thielscher et al., 2015) on this mesh, using the PARDISO direct solver and an isotropic conductivity assignment to skin, skull, cerebrospinal fluid, grey matter, and white matter. Anisotropic conductivity is not used because it injects a correlated bias into the calibration ratio; details in Section 2.11. Every simulation in the pipeline runs at a fixed reference current slew rate of d*I*/d*t* = 10^6^ A/s (1 A/μs), so that the resulting field magnitudes are intrinsic efficiencies in V/m per A/µs and can be rescaled to any stimulator intensity by a single multiplication.

The coil position is set by one of two routes: a user-supplied 4×4 ‘matsimnibs’ matrix, applied as given, or a cortical target coordinate, from which the pose is obtained by auxiliary dipole method optimisation (ADM, Gomez et al., 2021). The induced field is then sampled at every streamline point by tetrahedral interpolation on the FEM solution, in the shared RASMM space of Section 2.2, and passed to the AF computation of Section 2.4. The two pose-specification routes, the choice of optimiser method, the coil-to-scalp distance, the scalp projection with its default handle orientation, and the sampling routine are described in Supplementary Methods S7; the field-solver threading and the field-sample-matching safeguards that validate the sampled values against the requested streamline coordinates are described in Supplementary Methods S2.

### 2.6 Per-streamline contiguous-activation threshold

A single supra-threshold value of the activating function at one point on a fibre is not sufficient to evoke a propagated action potential: initiation requires the depolarising drive to be sustained over a span of membrane comparable to the internodal distance. We therefore reduce each streamline to a physically motivated scalar, defined as the largest activating-function magnitude that can be supported over a contiguous segment of pre-specified length.

Let (*AF_j_*) denote the signed AF values of Section 2.4, carried at the (M) midpoints of a streamline’s native segments, and let (*p_j_*)denote those midpoint coordinates. Restricting to the midpoints that fall inside the ROI leaves n ≤ M retained values. The threshold is evaluated on the intervals between consecutive retained midpoints: for (*j* = 1, …, *n* − 1) we take the local activating-function magnitude as the mean of the two adjacent absolute values, 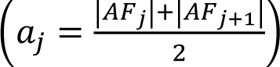, and the interval length as the Euclidean distance between the bounding midpoints, (*l_j_* = ‖*p_j_*_+1_ − *p_j_*‖), in millimetres. Magnitudes are averaged rather than the average taken in magnitude, so that a sign reversal between adjacent segments does not cancel to an artificially low local value. The per-streamline activation threshold is:

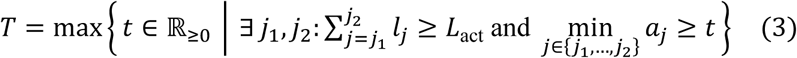

where *L*act is the user-configurable activation length (4 mm by default, set through ‘*options.activation_length_mm*’). *T* is therefore the largest activating-function magnitude such that at least one contiguous segment of total length not less than *L*act has all its midpoint values above *T*. A streamline that fails to deliver a contiguous supra-threshold segment of the required length is assigned *T* = 0.

The contiguous criterion makes the estimator robust to single-point AF spikes produced by residual numerical noise in the field gradient or by leftover tractography noise: an isolated spike on one segment cannot satisfy Eq. (3) by itself, so the streamline-level threshold is controlled by the geometry of an extended high-AF region rather than by the largest isolated value on the streamline.

The threshold is computed only on segments that lie inside the ROI sphere (Section 2.2). Outside the ROI the field has decayed, and the AF carries no information about activation at the targeted site. A streamline enters the bundle-level aggregate only if its retained portion supports the estimator: it must contribute at least two midpoints inside the ROI, all of its in-ROI AF values must be finite (Supplementary Methods S2), and its accumulated in-ROI path length must reach 10 mm. The last criterion excludes streamlines that clip the edge of the sphere over a span shorter than the shortest configurable activation length, for which Eq. (3) would be evaluated on too little support to be meaningful; it is a fixed internal constant rather than a user-facing option, and the streamline counts reported in the run log are the counts that survive it.

The default *L*act = 4 mm matches the order of magnitude of the activation length scale reported for myelinated cortico-cortical axons under standard TMS pulse parameters (Aberra et al., 2020). The activation length is exposed as options.activation_length_mm, so its effect on the bundle-level metric, the calibration ratio, and the inferred I_TIDE can be characterised directly on a user’s own data; the 2–8 mm range brackets the plausible internodal spacing of myelinated cortico-cortical axons at standard TMS pulse widths.

### 2.7 Cross-streamline aggregation

Equation (3) produces one threshold *Ti* per streamline that survives the four-point cutoff of Section 2.4. The bundle-level activating function used in the calibration ratio of Section 2.9 must aggregate these per-streamline thresholds into a single scalar. The choice of aggregator is not innocuous. A mean is biased downward by short or marginally activated streamlines, the maximum is biased upward by tractography artefacts that the angular filter of Section 2.4 does not catch, and a single high-order percentile has high variance when the streamline count inside the ROI is moderate. We adopt the median of the top five per cent of Ti as the bundle-level activating function and denote it AF_bundle_.

The aggregator is defined operationally as a two-step quantile composition. First, compute the 95th percentile of the per-streamline threshold distribution, *T*^(95)^ = *Q*0.95({*Ti*}), and retain the subset of streamlines for which *Ti* ≥ *Q*0.95({*Ti*}). Second, report the median of Ti over the retained subset:

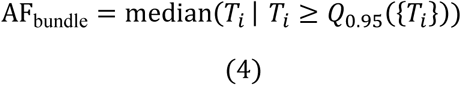

Two variants of AF_bundle_ are computed in parallel and reported side by side in every output. The unweighted variant treats every surviving streamline as equally informative and uses the empirical percentile and median. The weighted variant accepts an external per-streamline weight, by default a SIFT2 weight produced upstream by the tractography pipeline (Smith et al., 2015) and replaces the empirical percentile and median by their weight-aware counterparts in which each streamline contributes a fraction of the cumulative weight rather than a single count. The weighted variant is the physics-anchored estimate when SIFT2 weights are available, since the weighting compensates the per-streamline density bias of streamline-tracking algorithms; the unweighted variant is retained as a sensitivity reference. Where the two variants diverge, the interval they span is the appropriate expression of the uncertainty introduced by streamline density, and we recommend reporting both rather than selecting one.

The same primary aggregator is applied to the calibration bundle and to the target bundle. Identity of the aggregation rule on both sides of the calibration ratio used in Section 2.9 is required for the ratio to reflect a physical difference between the two bundles rather than an artefact of the aggregation choice. The median of the top 5% remains the only aggregator used to compute the reported dose. Five additional aggregators (median of the top 1%, Q0.95, Q0.90, median, and mean) are evaluated on the same per-streamline threshold distributions and reported as diagnostic sensitivity analyses; none alters the primary estimate or requires an additional field solve or activating-function computation. Their definitions and reporting are described in Supplementary Methods S8.

### 2.8 Weighting and surface-constraint modes

The unweighted aggregator of Eq. (4) is the safe default for tractograms without weight files; when SIFT2 weights are supplied, the weight-aware variant is reported and is the default for downstream consumers. Two further options, a grey-white surface constraint and a hybrid surface-plus-weight mode, together with the on-load validation of a supplied weight file and the streamline-index realignment that keeps each surviving streamline paired with its own weight after upstream drops, are described in Supplementary Methods S3.

### 2.9 Calibration anchor and target intensity inversion

The bundle-level activating function AF_bundle_ of Section 2.7 is an electric quantity expressed in V/m² per A/µs (intrinsic efficiency form, Section 2.5). To translate this quantity into a clinically actionable stimulator setting, an external anchor is required that maps an activating-function magnitude to a biological activation event. The pipeline uses the subject’s measured RMT on the M1 corticospinal tract as that anchor.

The calibration logic is the following. The RMT is, by definition, the stimulator intensity at which a defined population of corticospinal axons under the M1 hotspot fires reliably enough to generate a motor evoked potential of a fixed amplitude criterion in a target muscle (in our reference subject, the first dorsal interosseous; Rossini et al., 2015). At an intensity equal to RMT, the activating-function magnitude observed on the CST is therefore the magnitude that the subject’s CST axons require to reach their biological activation threshold. The pipeline samples this magnitude as AF_CST_, the cross-streamline aggregate of Section 2.7 evaluated on the CST tractogram at the coil position that produced the empirical RMT.

For any other target bundle, the stimulator intensity that produces the same activating-function magnitude AF_CST_ on the target bundle, and therefore the same biological activation event under the assumption that the biological threshold is approximately conserved across myelinated white-matter axons of comparable diameter, is given by the linear inversion

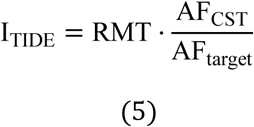

where I_TIDE and RMT denote percentages of the device maximum d*I*/d*t* and both AF_CST_ and AF_target_ are sampled at the same reference d*I*/d*t* = 1*x*10^6^ A/s (Section 2.5). The ratio is therefore dimensionless, and the inversion is linear in RMT.

Equation (5) admits an interpretation as a per-target dose multiplier. Setting

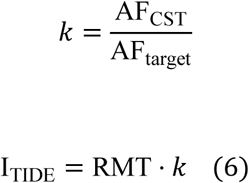

The multiplier *k* is a property of the subject’s anatomy and of the chosen coil poses and is independent of RMT. Once computed, *k* allows the user to rescale the predicted target dose to any RMT, including a later remeasurement on the same subject, without re-running the FEM simulation. The decoupling holds exactly under the linearity of the FEM solver and the choice of a common reference d*I*/d*t* for the CST and the target simulations.

The pipeline additionally reports a dimensionless figure of merit, the **Stimulation Efficiency Index (SEI)**, defined as the reciprocal of the multiplier:

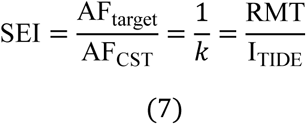

SEI quantifies how efficiently a target bundle is activated by a given coil pose, relative to the calibration bundle under the same subject and the same RMT measurement. SEI = 1 identifies the calibration condition (target activation parity with the CST), recovered exactly in the M1 identity case in which calibration and target are the same bundle. SEI > 1 identifies a target bundle that is more efficient than the CST under the chosen coil pose, so a lower stimulator output suffices to reach the biological threshold. SEI < 1 identifies a target bundle that is less efficient than the CST, so a higher stimulator output is required. The relative deviation of the inferred dose from the RMT admits a closed-form expression in SEI. From Eq. (7),

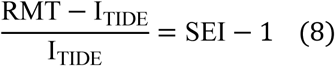

so, SEI-1 is the signed fractional difference between RMT and I_TIDE, taken relative to I_TIDE. To first order around the calibration condition this is also the fractional shift relative to RMT, and *SEI* − 1 provides a quick scalar summary of how far the target estimate departs from the motor threshold without consulting the full report. Because the numerator and denominator of Eq. (7) are two activating functions computed in the same subject, from the same simulation and at the same stimulator output, SEI is independent of the intensity anchor: rescaling RMT rescales I_TIDE by the same factor and leaves SEI unchanged. The relative deviation of Eq. (8) is therefore a measure of pathway geometry and not of delivered intensity, a property on which the validation design of Section 2.12 depends.

Weighted and unweighted variants of Eqs. (5)–(7) are computed in parallel and populate the corresponding rows of the results table.

Eq. (5) rests on two assumptions: the biological activation threshold of myelinated cortical axons is approximately conserved across bundles of comparable diameter and myelination, and the RMT has been measured under the same head model and M1 coil pose used in the simulation. A re-measurement is required if either changes.

Because Equation (5) is linear in the motor-referenced intensity, the multiplier can be reused to update after a new RMT measurement or a change in the protocol-specific RMT scaling factor, without repeating the FEM simulations. The raw estimate is not intrinsically restricted to the output range of the stimulator or to the intensity interval specified for a particular protocol. TIDE therefore reports both the raw estimate, ITIDE, and a bounded value, ITIDE,clamped, obtained by applying user-defined lower and upper reporting limits. The default limits are 0.70 x RMT and 1.40 x RMT, with the upper value additionally restricted by the maximum stimulator output of the device. Each estimate is assigned one of five status flags: WITHIN_RANGE when the raw estimate lies within the configured interval; CLAMPED_LOW or CLAMPED_HIGH when it falls below or above that interval; DEVICE_LIMITED when the requested intensity exceeds the device output range; and ESTIMATION_FAILED when the inversion cannot be evaluated. The last condition occurs when the target AF aggregate is zero, undefined, or non-finite, for example because no eligible streamline contributes to the estimate within the ROI. In this case, both the raw and bounded intensities are reported as missing and the observation is excluded from downstream summaries, thereby distinguishing an estimation failure from a valid low-intensity estimate. The reporting limits do not alter *k*, SEI, or any upstream AF quantity, and they should not be interpreted as physiological safety limits. The bounding equation and the relationships among the status categories, *k*, and SEI are provided in Supplementary Methods S9.

### 2.10 Operational Workflows

The numerical core described above is exposed through four operational workflows that share a common configuration schema and SimNIBS interface but differ in scope. Two workflows perform the calibration inversion of Eq. (5) on different scales: a single-pair estimation and a multi-position grid search over a cortical surface. Two utility workflows expose the underlying SimNIBS field solver and coil optimiser without performing the calibration inversion; the simulation workflow can additionally map the sampled field onto a supplied target bundle.

#### 2.10.1 Estimation workflow

The estimation workflow (*tide --workflow estimation*) processes one calibration bundle and one target bundle in a single run. The unified estimation block then evaluates Eqs. (5)–(8), applies the intensity clamp (Section 2.9; Supplementary Methods S9), and reports weighted and unweighted I_TIDE in both raw and clamped form, together with an independent clamp flag for each variant. The report also includes the aggregator-sensitivity diagnostics described in Section 2.7; these do not alter the primary dose estimate.

A self-validation block re-samples the cached target field on the M1 mesh to test calibration–target consistency at no additional FEM cost; the report fields it emits are described in Supplementary Methods S4.

The output directory layout, the report-block order, and the companion artefacts are listed in Supplementary Methods S4. An automatically optimised coil pose that fails the coil-pose plausibility check of Supplementary Methods S4 is rejected before the calibration inversion, so no intensity is reported for an implausible skull-base pose.

#### 2.10.2 Grid-search workflow

The grid-search workflow (*tide --workflow grid*) evaluates the calibration inversion at many candidate cortical positions on the target bundle and emits separate scalar maps of the raw and clamped intensity, together with a categorical map of the clamp regime, over the cortical surface. The candidate set is built from the union of the streamline start and end points, reduced in one of two ways. When the user supplies a seed cortical coordinate, the endpoints are restricted to a sphere of radius grid.search_radius_mm around it, which is the usual mode of operation. When no seed is supplied, the pooled endpoints are instead partitioned into two clusters by K-means on the tip coordinates and the superior cluster is retained, on the assumption that the bundle has one cortical and one subcortical or contralateral termination. The two reductions are alternatives, not successive filters. The surviving endpoints are then restricted to a superficial band, defined as the interval of width grid.cortex_depth_mm below the 98th percentile of the endpoint z coordinates, and quantised to a configurable lattice step (grid.step_size_mm, 4 mm by default) with duplicates removed, so that candidate positions are commensurate across runs. The depth criterion is expressed on the scanner z axis rather than on the local scalp normal, which is appropriate for dorsal and superior targets and increasingly conservative for lateral and inferior ones.

The run-wide artefacts, the per-point outputs that mirror a single estimation run, and the cortical-surface visualisations are listed in Supplementary Methods S4.

#### 2.10.3 Standard simulation and optimisation

Two utility workflows expose the underlying SimNIBS components without performing the calibration inversion. *tide --workflow simulation* runs a single FEM solve at a user-specified coil pose and, when a target bundle is supplied, can additionally map the E-field or activating function onto that bundle. *tide --workflow optimization* runs the SimNIBS coil-position optimiser at a user-specified cortical target; an optional tractogram can be used to derive a cortical medoid target. Neither workflow evaluates Eq. (5), so neither produces a TIDE_Results_<target>.txt summary.

The outputs are correspondingly minimal when no tractogram is supplied. The standalone simulation writes the SimNIBS simulation directory and the YAML configuration snapshot; when a target bundle is supplied, it also writes the mapped streamline scalars and associated analysis artefacts. The standalone optimisation writes the 4×4 matsimnibs matrix, the scalp coordinate, and the YAML configuration snapshot. Either workflow is appropriate for users who require the induced E-field at a fixed pose, tract-level field mapping without dose inversion, or an optimised coil pose as input to a separate downstream analysis.

### 2.11 Design: conductivity model and relation to E-field-only optimisation

The finite-element solver of Section 2.5 assigns scalar (isotropic) conductivities to the five anatomical tissue tags of the head mesh and does not accept the diffusion-tensor-derived anisotropic conductivity that SimNIBS exposes as an optional configuration. This is a deliberate design choice rather than a limitation of the implementation. The decoupling of the tractogram (the geometric carrier of the activating function) from the volume conductor (the electric carrier of the field) is what makes the calibration ratio of Eq. (5) interpretable as a comparison of bundle geometries under a common electrical model. An anisotropic FEM conductivity derived from the same diffusion-MRI dataset that produced the tractogram would inject a correlated bias into both terms of that ratio, and the calibration would lose its anatomical specificity.

An anisotropic tensor built from the same diffusion data that produced the tractogram correlates with fibre orientation by construction, so it amplifies the activating-function ratio AF CST/AF target unequally on the two bundles and makes the inferred *I*_*TIDE* depend on the DTI-to-conductivity mapping; the mechanism, and the E-field-modelling literature that supports the choice on independent grounds, are given in Supplementary Methods S6.

The AF magnitudes reported by the pipeline therefore depend on the isotropic SimNIBS conductivity assignments (Saturnino et al., 2019), while the geometric content of the bundle is carried independently by the Frenet–Serret frame of Section 2.4. The tractography pathway and the electric pathway are independent inputs to the calibration ratio, which is the desired property for the cross-bundle comparison.

A further reference point is the E-field optimisation literature that informs the cortical hotspot selection of the conventional protocol (Gomez et al., 2021; Saturnino et al., 2019). This literature solves the problem of where to place the coil so that the cortical E-field magnitude at the user-specified target is maximised. TIDE uses the same SimNIBS optimiser (ADM, Section 2.5) for the same purpose. The E-field optimisation is therefore an upstream component of TIDE, not a competing tool: it produces the coil pose that the activating-function analysis then evaluates. The contribution of TIDE is the post-optimisation translation of the E-field into a target-specific stimulator output via the calibration ratio. Users who require only the coil pose and the surface E-field, for example for transcranial electrical stimulation or for an analysis that does not consume a tractogram, are directed to the standalone simulation and optimisation workflows of Section 2.10.

### 2.12 Validation against an independent behavioural cohort

The calibration arithmetic of Eq. (5) translates a per-bundle activating-function magnitude into a stimulator setting under the working assumption that the biological activation threshold is approximately conserved across myelinated cortico-cortical axons of comparable diameter and myelination (Section 2.9). We tested whether the cross-bundle dose differences produced by this inversion carry information about cortex-level behavioural variance by applying the pipeline to the data of (Tagliaferri et al., 2023). In that study, 19 healthy adults received single-pulse TMS delivered during the set-period of a precued reaction-time task at the six cortical endpoints of the left FAT (P01–P06: posterior, middle, and anterior sub-bundle terminations on the superior and inferior frontal gyri). The primary readout is a reactivity index (RI), bounded between 0 (fully predictive) and 1 (fully reactive). The published behavioural effects are confined to two endpoints: stimulation of the middle SFG site (P03) shifted choices toward predictive behaviour, and stimulation of the middle IFG site (P04) shifted choices toward reactive behaviour. The four remaining endpoints produced no detectable behavioural shift.

The validation tests whether the per-subject discrepancy between the experimentally delivered intensity I_delivered and the intensity I_TIDE recommended by the inversion of Eq. (5) predicts the per-subject behavioural response at each of the two effect endpoints. The dose recorded in the original protocol is RMT × 120%. The discrepancy metric is the unsigned relative deviation

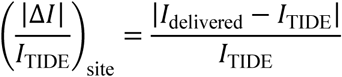

evaluated on the raw output of Eq. (5) before the clamp (Section 2.9; Supplementary Methods S9) is applied; the clamp is a reporting constraint and a wet-lab safety bound (Section 2.9) and is not propagated to inferential tests. The behavioural readout is the sham-corrected shift in the reactivity index, with a gyrus-specific sign chosen so that a single one-tailed hypothesis applies to both arms. At the SFG site, the shift is Δ*RIP*03 = *RI*sham − *RIP*03, positive when TMS pushed the subject toward predictive choices. At the IFG site, the shift is Δ*RIP*04 = *RIP*04 − *RI*sham, positive when TMS pushed the subject toward reactive choices. The predicted direction of the dose–behaviour coupling is the same under both definitions: a smaller |ΔI| / I_TIDE, i.e. a delivered intensity closer to the recommendation, should associate with a larger behavioural shift along the gyrus-appropriate axis. The pre-registered statistical test is Spearman’s rank correlation ρ(|ΔI| / I_TIDE, ΔRI), one-tailed at α = 0.05, with a 95% percentile bootstrap confidence interval on ρ over 10 000 resamples. Because the delivered dose is itself the intensity anchor supplied to Eq. (5), the tested predictor reduces by Eq. (8) to |ΔI| / I_TIDE = |SEI − 1 |: the anchor cancels between numerator and denominator, and the predictor is a function of the AF_target / AF_CST ratio alone. The magnitude of the delivered intensity cannot enter the correlation, so a monotone relationship between stimulator output and behavioural shift could not by itself generate a non-zero ρ. Two confound tests were specified in advance so that this property is demonstrated in the data and not only in the algebra: ρ(I_delivered, ΔRI), which asks whether the delivered intensity predicts the behavioural shift directly, and ρ(I_delivered, |ΔI| / I_TIDE), which asks whether the delivered intensity determines the discrepancy metric. We further pre-specified that the signed recommendation I_TIDE − I_delivered would be reported for each of the six protocol sites, because a predictor taken in absolute value conceals whether the pipeline recommends an increase or a reduction.

The TIDE estimation for the validation used each subject’s measured RMT × 120% as the calibration anchor (Section 2.9), the subject’s TractSeg-reconstructed FAT bundle (Section 2.3), the cortical target coordinates of the original protocol, and isotropic FEM conductivity (Section 2.11). All cohort-level analysis scripts, the pre-registration record, and the table of per-subject discrepancies are released alongside the pipeline at the Zenodo archive accompanying this manuscript (10.5281/zenodo.22019794).

As a secondary analysis (**Figure 3**) of the shape of the dose-behaviour relationship, we retained the sign of the TIDE recommendation and defined the signed relative dose adjustment as 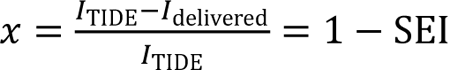. Thus, negative values indicate that TIDE recommends a lower intensity than was delivered, and positive values indicate that TIDE recommends a higher intensity. For each effect site separately, we fitted the unconstrained quadratic model ΔRI = *β*_0_ + *β*_1_*x* + *β*_2_*x*^2^ + *ε*. The point *x* = 0 was used as a physiological reference, so when *β*2 < 0 the location of the quadratic maximum was estimated as 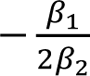. Ninety-five per cent confidence bands for the fitted curve were obtained by percentile bootstrap over participants with 10,000 resamples.

**Figure 3.**
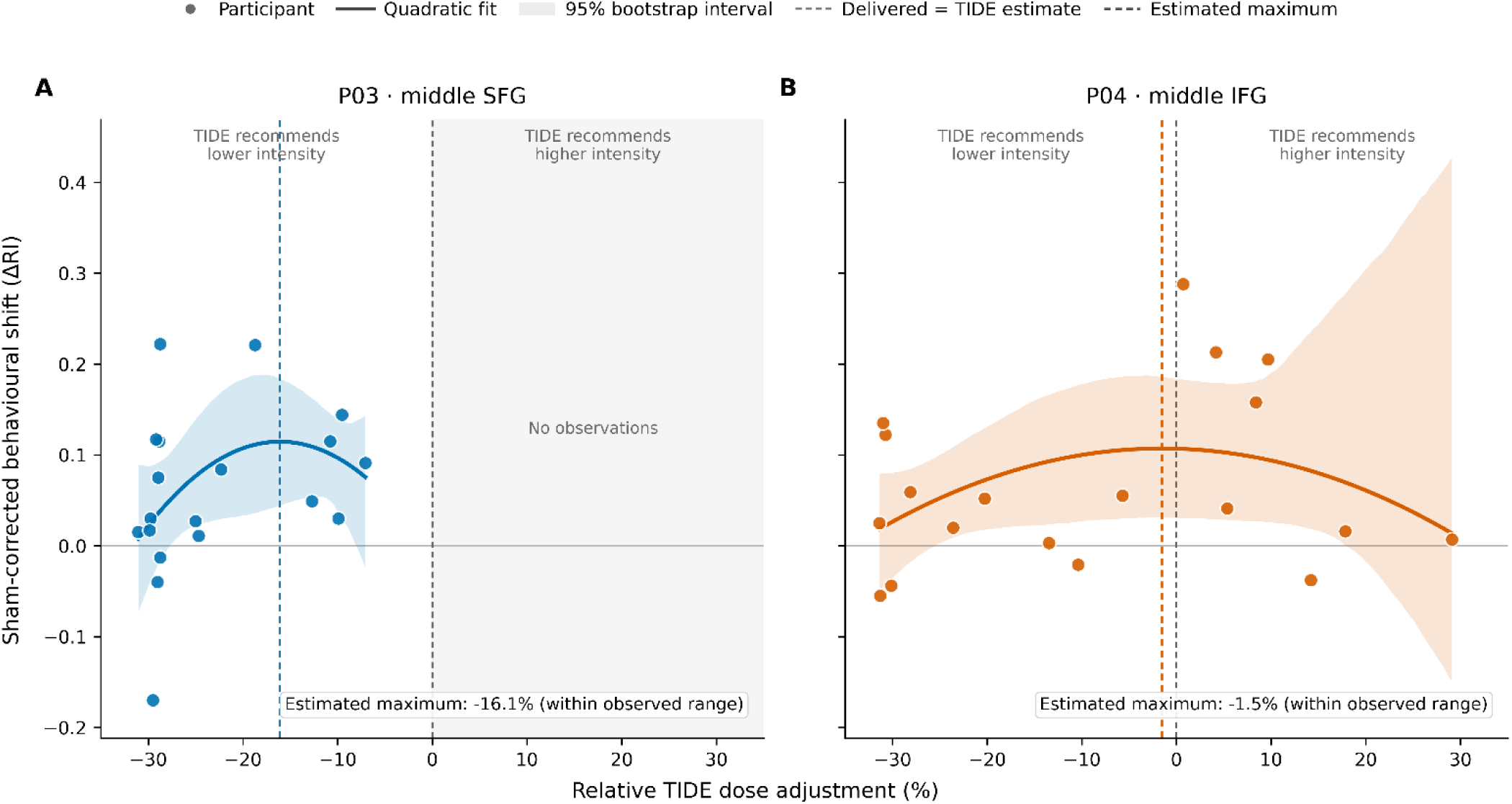
Quadratic relationship between relative TIDE dose adjustment and behavioural response at the two FAT effect sites. Sham-corrected behavioural shifts are plotted against the signed relative TIDE dose adjustment, defined as (*ITIDE* − *Idelivered*)/*ITIDE* and expressed as a percentage. Positive behavioural values denote the site-specific effect observed in the original study: a shift towards predictive behaviour at P03 and towards reactive behaviour at P04. The grey dashed line at 0% marks equality between the delivered and TIDE-estimated intensities; negative values indicate that TIDE recommends a lower intensity than was delivered, whereas positive values indicate that TIDE recommends a higher intensity. Solid curves show unconstrained quadratic fits and shaded areas show 95% participant-bootstrap confidence intervals. Coloured dashed lines mark the maxima estimated from the quadratic fits. **(A)** At P03 (middle SFG), all participants were on the negative side of dose matching and the estimated maximum was −16.1%. **(B)** At P04 (middle IFG), observations spanned both sides of dose matching and the estimated maximum was −1.5%, close to the point of agreement between delivered and TIDE-estimated intensity. The 0% reference was not imposed on either fit. SFG, superior frontal gyrus; IFG, inferior frontal gyrus; RI, reactivity index.

We also performed two secondary random-assignment analyses to test whether the observed subject-specific TIDE estimates carried more behavioural information than arbitrary assignments of the same estimates across participants. The first analysis permuted the absolute I_TIDE %MSO, whereas the second permuted the Stimulation Efficiency Index (SEI) and therefore removed the individual absolute intensity scale from the randomised predictor. Behavioural outcomes and delivered intensities were held fixed in both analyses. The permutation procedures and their distinct null hypotheses are described in Supplementary Methods S10.

### 2.13 Reference implementation, performance, and reproducibility

The pipeline supports Python 3.9–3.12 and is distributed on PyPI as the installable package tide-pipeline (command-line entry point: *tide*; Python import package: *tide*) under the GNU General Public License v3.0 or later (GPL-3.0-or-later). The software version associated with this manuscript is TIDE v1.30.0, archived on Zenodo under DOI <u>10.5281/zenodo.22019737</u>. A canonical annotated configuration can be generated directly from the installed package with *tide --init-config config.yml*. The FEM solve and the ADM coil optimisation are deterministic under fixed inputs: repeated runs of the same configuration reproduce the coil matsimnibs pose and the downstream AF and I_TIDE to full precision within the pinned software environment and stated numerical tolerances. PyPI packaging and distribution, platform support, environment bootstrapping, the command-line entry point and versioning scheme, the content-addressed fixed-pose cache, and the regression and sanity-check test suite are described in Supplementary Methods S5.

## 3. Results

### 3.1 Pipeline-recommended dose tracks behavioural variance at the FAT effect sites

We applied the pipeline to the 19 participants from Tagliaferri et al., (2023), who performed a cued reaction-time task during single-pulse TMS. For each subject we compared the delivered intensity — the lab dose, I_delivered = RMT × 120 % — with the intensity the pipeline recommends for that target (I_TIDE, the calibration output defined in Methods, Eq. 5), and took their relative deviation 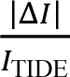. We evaluated this at the two FAT endpoints that produced a behavioural effect in the original study: the middle superior-frontal-gyrus site (SFG; P03) and the middle inferior-frontal-gyrus site (IFG; P04).

The cohort-level behavioural shifts were reproduced in our re-analysis. P03 stimulation moved 16 of 19 subjects toward predictive choices, reflected by shorter reaction times (one-sample Wilcoxon signed-rank test, p = 0.005), and P04 stimulation moved 15 of 19 toward reactive choices, reflected by longer reaction times (Wilcoxon, p = 0.011). The two shifts load on opposite poles of the reactivity index (RI), which matches the published dissociation between the superior and inferior frontal terminations of the middle FAT sub-bundle. The two site-specific correlations were the pre-specified tests (Methods, Section 2.12).

The recommendation was predominantly downward. At P03 the pipeline recommended a lower intensity than the delivered RMT × 120 % dose in all 19 subjects (median I_TIDE 42.0% % of maximum stimulator output against a delivered 54.0 %, a median reduction of 22.2 %), and the same direction held at the two other superior-frontal sites (P05, 17 of 19; P01, 16 of 19). The inferior-frontal sites divided in both directions (P04, 11 of 19 downward; P02 and P06, 9 of 19 each). Across the six protocol sites and 19 subjects, 81 of 114 recommendations fell below the delivered dose, and the physiological clamp of Section 2.9 was engaged 38 times at its lower bound and never at its upper bound. The association reported below is therefore not a restatement of the proposition that a larger stimulator output yields a larger effect: the pipeline recommended a larger output in a minority of observations and, at the site carrying the superior-frontal arm, in none of them (Table 1).

**Table 1.** Direction of the TIDE recommendation relative to the delivered dose at the six protocol sites. For each stimulation site of Tagliaferri et al. (2023), the number of participants for whom the pipeline recommended an intensity below or above the delivered RMT x 120 % dose, the median recommended intensity, and the median relative change with respect to the delivered dose. Values are the SIFT2-weighted aggregate; n = 19 per site. The delivered dose had a median of 54.0 % of maximum stimulator output (range 37 to 67). Across all 114 site-participant observations the physiological clamp of Section 2.9 was engaged 38 times at its lower bound and never at its upper bound. SFG, superior frontal gyrus; IFG, inferior frontal gyrus.

| Site | Gyrus | $I_{\text{TIDE}}$ below delivered | $I_{\text{TIDE}}$ above delivered | Median $I_{\text{TIDE}}$ (% max output) | Median relative change |
| --- | --- | --- | --- | --- | --- |
| P01 | SFG | 16 | 3 | 43.2 | −9.3 % |
| P02 | IFG | 9 | 10 | 50.9 | +0.4 % |
| P03 | SFG | 19 | 0 | 42.0 | −22.2 % |
| P04 | IFG | 11 | 8 | 48.6 | −9.4 % |
| P05 | SFG | 17 | 2 | 44.4 | −19.1 % |
| P06 | IFG | 9 | 10 | 50.8 | +0.3 % |

Because both tested the same hypothesis—that a delivered dose closer to the TIDE recommendation would be associated with a larger behavioural effect in the direction observed at each site—we also combined the two sites in a single pooled analysis. To express the behavioural effect on a common scale, we defined the sham-corrected shift so that positive values always indicated movement in the previously observed site-specific direction. The shift was calculated as RI_sham - RI_P03 for the SFG site and RI_P04 - RI_sham for the IFG site. Each of the 19 participants therefore contributed two observations, one for each site. For each observation, we considered the relative discrepancy between the delivered and recommended dose, 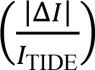, and the corresponding sham-corrected behavioural shift. Pooling the 38 observations yielded a Spearman correlation in the predicted negative direction (ρ = −0.39), indicating that smaller deviations from the TIDE-recommended dose were associated with larger behavioural shifts. Because the two observations contributed by each participant were not independent, significance was assessed using a subject-clustered permutation test in which each participant’s two-site block was permuted as a single unit (10,000 permutations). The confidence interval for ρ was estimated using a subject-clustered bootstrap. The pooled association was significant (one-tailed permutation p = 0.012; bootstrap 95% CI [−0.70, −0.02]). As the pooled analysis represents a single combined endpoint, no multiple-comparison correction was applied to this test. A conventional Spearman test that ignored the repeated-measures structure yielded p = 0.008, indicating that accounting for the two observations per participant did not inflate the significance of the association.

The site/gyrus-specific analyses yielded associations of the same direction and magnitude. At P03 the dose discrepancy and the predictive shift are rank-correlated in the predicted direction (Spearman ρ = −0.44, one-tailed p = 0.028, 95% bootstrap CI [−0.71, −0.05]; n = 19). At P04 the reactive shift gives a correlation of the same sign and size (ρ = −0.44, one-tailed p = 0.029, 95% CI [−0.79, +0.10]; n = 19). Read as two co-equal tests, they call for a multiple-comparison correction. Under the pre-specified Benjamini–Hochberg false-discovery control, the result holds across both arms (adjusted p = 0.029 at both sites). As a stricter, more conservative check, the finding does not clear family-wise control (Bonferroni-Holm adjusted p = 0.057 and p = 0.058). The two arms fall just inside the false-discovery threshold and just outside the family-wise threshold, which is the expected position for two positively correlated tests whose uncorrected p-values sit near α/2. We report the uncorrected, false-discovery, and family-wise results together rather than only the most favourable one. The two pre-specified confound tests (Section 2.12) returned null associations. Delivered intensity did not predict the behavioural shift at either site (P03, ρ = +0.08, p = 0.76; P04, ρ = −0.06, p = 0.82; pooled across the 38 observations, ρ = +0.03, p = 0.86), and it did not determine the discrepancy metric (P03, ρ = +0.28, p = 0.25; P04, ρ = +0.20, p = 0.40). At P04, where participants occurred on both sides of the TIDE recommendation, the association was present among the eight subjects whose delivered dose lay below the recommendation (ρ = −0.83, p = 0.010), but not among the eleven dosed above it (ρ = +0.05, p = 0.87). Within that under-dosed subgroup the delivered intensity is not the variable that moves. The spread in |ΔI| / I_TIDE originates in I_TIDE, that is in bundle geometry, so a larger recommended intensity accompanies a smaller behavioural shift at an essentially fixed delivered dose.

The unsigned discrepancy used in the primary analysis does not distinguish whether the delivered intensity lay above or below the TIDE estimate. We therefore examined the signed relationship in a quadratic analysis (**Figure 3**). At P04, where observations occurred on both sides of dose matching, the freely estimated maximum of the quadratic curve was at a relative TIDE dose adjustment of −1.5%, close to the point at which the delivered and TIDE-estimated intensities coincide. At P03, the estimated maximum was at −16.1%, within the observed range, but all participants remained on the same side of the TIDE estimate. The P04 pattern is therefore compatible with a behavioural maximum near dose matching, whereas the P03 data cannot test a two-sided maximum around 0%.

We next tested whether the observed subject-specific mapping between TIDE estimates and behavioural responses was more informative than random reassignment of the same estimates across participants (Supplementary Methods S10). When the absolute TIDE intensities in %MSO were randomly reassigned while the observed behavioural responses and delivered intensities remained fixed, the true assignment produced a mismatch-behaviour correlation in the negative tail of the permutation distribution at both sites (P03: *ρ* = −0.444, *p* = 0.0018; P04: *ρ* = −0.442, *p* = 0.0352; Supplementary Figure S1). Because permutation of absolute %MSO also disrupts the association between the TIDE estimate and each participant’s individual stimulation scale, we repeated the test by randomising SEI directly. The observed correlations again fell in the negative tail of the corresponding null distributions (P03: *ρ* = −0.444, *p* = 0.0283; P04: *ρ* = −0.442, *p* = 0.0292; Supplementary Figures S2). Thus, the association was not restricted to the absolute intensity scale: the subject-specific target-to-CST efficiency estimated by TIDE also carried behavioural information that was generally lost when SEI values were reassigned across participants.

Across the pooled and site-specific analyses, smaller discrepancies between the delivered RMT × 120% dose and the TIDE estimate were associated with larger TMS-induced behavioural shifts along both the predictive SFG and reactive IFG axes of the original task. The shape analysis at P04 was compatible with a behavioural maximum close to dose matching, and the random-assignment analyses showed that the observed subject-specific TIDE and SEI mappings were uncommon under their respective permutation nulls.

Taken together, these retrospective results indicate that subject-specific TIDE estimates contain behaviourally relevant information beyond the delivered intensity alone.

### 3.2 Bundle geometry drives recommended-dose variance across the lateral association tracts

We applied the grid-search workflow to a cohort of 27 subjects with high-quality diffusion, each carrying the left corticospinal tract as the RMT-calibration anchor and thirteen target segments inside the figure-of-eight depth envelope: the arcuate fasciculus and the three superior longitudinal fascicles, each split into anterior and posterior cortical terminations; the inferior longitudinal and inferior fronto-occipital fasciculi; a premotor thalamo-cortical projection; and a parieto-occipital thalamic projection split at its parietal and occipital termini. Every bundle was reconstructed with the protocol of Supplementary Methods S1 (summarised in Section 2.3). TractSeg (Wasserthal et al., 2018) supplied the bundle mask, the endpoint regions, and the tract-orientation maps; probabilistic tracking inside the mask, seeded against the orientation prior, produced the streamlines; and SIFT2 (Smith et al., 2015) generated the per-streamline weights consumed by the weighted aggregator. The calibration corticospinal tract was reconstructed under the identical protocol, so anchor and target bundles enter the calibration ratio through the same tractography pathway. For every subject and bundle, the workflow searched scalp position and handle orientation over a 40 mm radius at a 6 mm step and returned the intensity that brought the bundle’s activating function up to the subject’s calibration value — the activating function on the corticospinal tract at RMT (*AF*_*CST*). This grid search produced 351 runs over 6,492 coil poses. The representative intensity for a (subject, bundle) pair was taken as the median of *I*_*TIDE* across that run’s grid points, and the per-bundle statistics aggregate the 27 per-subject medians.

One provenance fact bounds the interpretation of this interim run. The calibration RMT was held at 50 % max output for every subject rather than each subject’s measured threshold, so the between-subject spread reported below is geometric in origin (bundle course, cortical-endpoint alignment, scalp depth) and carries none of the real motor-threshold variance that a completed cohort will add. The Stimulation Efficiency Index (SEI; Methods Section 2.9) is immune to the placeholder, because SEI = RMT / I_TIDE cancels the shared anchor; SEI is therefore the transferable quantity here, and each subject’s I_TIDE will rescale by RMT_subject / 50 once real thresholds are supplied. The clamp window for these runs was 40–100 % max output, obtained with a floor ratio of 0.8 at the 50 % placeholder RMT; it is the window referenced throughout this section and is a per-run configuration choice, independent of the software defaults of Section 2.9.

The reference table this analysis set out to build proved defensible for none of the thirteen bundles [**Table 2** — per-bundle recommended intensity, SEI, and inter-subject variance]. Under a graded variance rule (coefficient of variation ≤ 15 % publishes a mean, 15–20 % is borderline, above 20 % calls for individualised estimation), no bundle cleared the publish-mean threshold: the lowest-variance target, T\_PREM\_left, fell just inside the borderline band (coefficient of variation CoV 15.8 %, *I*_*TIDE* 55.0 ± 8.7 % max output, 96 % of values inside the 40–100 % programmable window), and four further targets were borderline (SLF\_I\_left\_P 17.0 %, SLF\_II\_left\_A 17.9 %, *SLF*_*I*_*left*_*A* 19.1 %, *SLF*_*II*_*left*_*P* 19.3 %). No bundle reached a coefficient of variation below 15 %. Pooled across all 351 runs, the recommended intensity averaged 52.4 % max output at a median SEI of 1.00, with 77 % of the representative values inside the programmable window. The decision rule thus resolved to individualised estimation for eight of the thirteen bundles, with the remaining five borderline and none qualifying for a published group mean. That verdict is itself the result the pipeline is built to produce: geometry-aware dosing on lateral association tracts does not collapse to one number per tract, and the inter-subject variance is a measurement of that fact rather than a shortfall of the method.

**Table 2.** Per-bundle recommended intensity, Stimulation Efficiency Index (SEI), and inter-subject variance across the thirteen target white-matter bundles under the grid-search workflow (N = 27 subjects, 351 runs, 6,492 coil poses). Values summarise the recommended intensity (I_TIDE, % of maximum stimulator output) as mean ± SD, median, and interquartile range (IQR); sCoV, coefficient of variation; SEI (med), median Stimulation Efficiency Index; Window %, percentage of representative values inside the 40–100 % programmable window. Verdict follows the graded variance rule of Section 3.2 (CoV ≤ 15 %, publish mean; 15–20 %, borderline; >20 %, individualise). Bundles are ordered by increasing CoV.

| Bundle | Mean ± SD | Median | IQR | CoV % | SEI (med) | Window % | Verdict |
| --- | --- | --- | --- | --- | --- | --- | --- |
| T_PREM_left | 55.0 ± 8.7 | 54.4 | 48.7–62.6 | 15.8 | 0.92 | 96 | Borderline |
| SLF_I_left_P | 54.9 ± 9.3 | 52.6 | 47.7–63.2 | 17.0 | 0.95 | 96 | Borderline |
| SLF_II_left_A | 49.4 ± 8.9 | 49.8 | 42.9–56.6 | 17.9 | 1.00 | 78 | Borderline |
| SLF_I_left_A | 45.1 ± 8.6 | 44.1 | 40.3–48.3 | 19.1 | 1.13 | 74 | Borderline |
| SLF_II_left_P | 57.7 ± 11.2 | 56.3 | 49.0–66.7 | 19.3 | 0.89 | 96 | Borderline |
| AF_left_P | 47.5 ± 9.9 | 47.2 | 39.3–53.7 | 20.7 | 1.06 | 67 | Individualise |
| T_PAR_left_par | 55.0 ± 12.1 | 55.6 | 46.2–60.4 | 22.1 | 0.90 | 89 | Individualise |
| SLF_III_left_A | 50.3 ± 11.7 | 51.4 | 42.3–57.8 | 23.3 | 0.97 | 78 | Individualise |
| IFO_left | 34.2 ± 8.0 | 34.4 | 28.0–39.5 | 23.5 | 1.45 | 22 | Individualise |
| ILF_left | 49.3 ± 11.7 | 51.0 | 40.5–56.6 | 23.8 | 0.98 | 81 | Individualise |
| T_PAR_left_occ | 82.6 ± 20.5 | 78.6 | 70.1–90.6 | 24.8 | 0.64 | 85 | Individualise |
| AF_left_A | 55.8 ± 14.6 | 53.7 | 44.0–62.6 | 26.1 | 0.93 | 85 | Individualise |
| SLF_III_left_P | 44.4 ± 14.9 | 42.0 | 35.3–48.6 | 33.5 | 1.19 | 59 | Individualise |

The SEI ordering separated favourable from unfavourable bundle geometry. Four bundles activated more efficiently than the calibration CST (SEI > 1), led by IFO_left at 1.45 and followed by SLF_III_left_P (1.19), SLF_I_left_A (1.13), and AF_left_P (1.06); each carries a superficial, well-aligned cortical segment that the conventional RMT × 120 % convention would over-drive. Among the under-efficient bundles (SEI < 1), the clearest examples were the deep or poorly oriented terminations T_PAR_left_occ at 0.64, SLF_II_left_P at 0.89, and AF_left_A at 0.93, each of which required a higher output than the motor-referenced dose to reach the same activation.

**Figure 4** illustrates this contrast in a representative subject. Relative to the CST calibration bundle, AF_left_A and SLF_I_left_A follow distinct trajectories and cortical termination patterns, producing different spatial distributions of the activating function. These images are illustrative; the intensity and SEI estimates reported in Table 2 were derived from all 27 subjects and all eligible grid positions.

**Figure 4.**
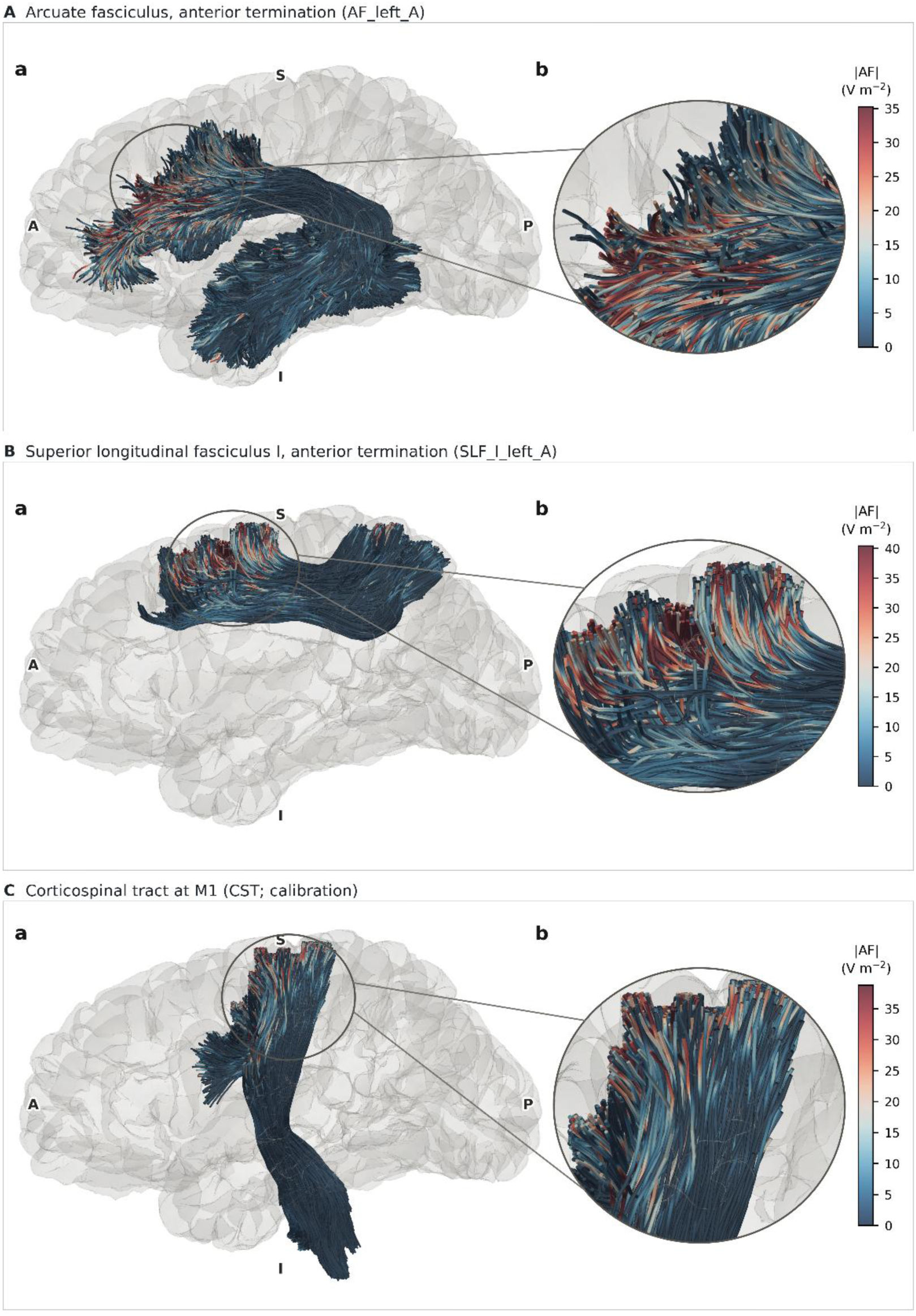
Representative tract geometry and activating-function distribution in the calibration and target bundles. **(A)** Anterior cortical termination segment of the left arcuate fasciculus (AF_left_A). **(B)** Anterior cortical termination segment of the left superior longitudinal fasciculus I (SLF_I_left_A). **(C)** Left corticospinal tract beneath M1 (CST; calibration). Within each row, subpanel (a) shows the complete tractogram in a lateral view of the left hemisphere, and subpanel (b) enlarges the circled cortical termination region. A, P, S and I denote anterior, posterior, superior and inferior, respectively. Colour represents the activating function in V m⁻² at the reference simulation rate of dI/dt = 1 A/µs. The display limit was set independently for each bundle at the 99th percentile of |AF| to reduce the influence of extreme values; colours should therefore be interpreted within rows and should not be compared quantitatively across rows. All three tractograms contain 25,000 streamlines and are shown for a representative subject. Cohort-level intensity and SEI estimates are reported in Table 2.

Two bundles sat at the boundary of deliverable stimulation and warrant an explicit depth-limited caveat rather than a recommended mean. T_PAR_left_occ, a deep parieto-occipital thalamic projection, had the highest mean intensity at 82.6 % max output, reached 146 % in the most extreme subject, and produced every above-ceiling value in the pooled set, placing it at or beyond what a figure-of-eight C-B60 can drive. IFO_left failed in the opposite sense: its SEI of 1.45 pushed the recommended intensity down to a 34.2 % mean, leaving only 22 % of subjects inside the programmable window, so most recommendations fell below the 40 % floor. Both tracts lie at the edge of the depth envelope the bundle selection was meant to enforce.

Clamping at the individual grid-point level was common but expected. Of the 6,492 poses, 68 % were within range, 24 % clamped low, and 8 % clamped high, and no run produced a non-finite activating function. Because these counts span the full searched lattice, including deliberately sub-optimal poses, they overstate clamping at the recommended pose, where the per-subject median lay within range for most bundles. Coil-pose scatter across subjects was not computed here, because native-space scalp coordinates are not comparable across subjects without the normalisation step reserved for a subsequent analysis; the reference table therefore reports intensity and efficiency but defers the geometric dispersion of the recommended pose.

## 4. Discussion

Two findings emerge from the present work. First, in an independent cohort, participants who received an intensity closer to the TIDE recommendation showed larger site-specific behavioural shifts, at both FAT endpoints at which the original study had reported an effect. Second, across thirteen white-matter targets, no bundle yielded a generalisable group-level dose: five were borderline and the remaining eight required subject-specific estimation. We consider each in turn.

The behavioural association supports the relevance of tract-informed dosing for experimental applications of TMS. It suggests that TIDE captured inter-individual differences in stimulation dose that were relevant to the behavioural consequences of targeting the frontal aslant tract (Tagliaferri et al., 2023). Importantly, larger behavioural effects were associated with smaller discrepancies between the delivered intensity and the TIDE recommendation, not with higher stimulation intensities per se. Consistently, the predictor was invariant to the intensity anchor, and the delivered intensity itself was not associated with the behavioural shift. Moreover, the pipeline recommended a reduction in stimulation intensity in 81 of 114 site–subject observations (across all stimulation sites in the source dataset), including every subject at the superior-frontal site. These findings argue against a simplistic explanation whereby larger stimulator outputs simply produce larger behavioural effects. Under such an account, stronger effects should have been associated with higher delivered intensities, and any improvement over the standard protocol should have resulted from prescribing more current. Neither pattern was observed. Although the present analysis does not establish that prospective delivery of I_TIDE would causally maximise the behavioural response, it provides evidence that improving tract-specific dose matching may enhance effective pathway engagement in a manner that translates into measurable behavioural effects.

The secondary analyses add two observations to this result. At P04, the maximum of an unconstrained quadratic fit fell close to the point at which delivered and TIDE-estimated intensities were equal. In addition, both the absolute-dose and SEI randomisation tests placed the true subject-specific assignments in the tails of their respective null distributions. The SEI analysis is particularly informative because the absolute RMT/%MSO scale cancels from the mismatch metric, leaving the target-to-CST efficiency ratio as the randomised quantity. These findings strengthen the evidence that the individual TIDE estimate contains behaviourally relevant information, but they do not establish I_TIDE as a causal optimum. A prospective experiment in which intensity is varied around I_TIDE within participants is required for that inference.

These findings also provide empirical support for the limitations of directly transferring an RMT-based intensity from the motor system to non-motor pathways. In the multi-bundle analysis, substantial between-subject variability in TIDE estimates emerged despite using the same placeholder calibration RMT (50% of maximum stimulator output) for all participants. The observed spread therefore reflected differences in pathway geometry, cortical-endpoint alignment and scalp depth, rather than variability in the MT anchor itself. For most examined bundles, these anatomical differences prevented tract-specific intensities from being reduced to a stable group-level value, supporting the need for subject-specific dosing. This result extends previous E-field modelling studies showing substantial inter-individual variation in the intensity required to reproduce a motor-equivalent field at non-motor targets (Caulfield et al., 2021; Numssen et al., 2024). TIDE retains the physiological information provided by RMT while translating it through the subject-specific geometry of the pathway intended to be engaged. Under conventional dosing, these anatomical differences remain uncontrolled sources of variance, whereas TIDE explicitly incorporates them into the dose estimate.

The response induced by a TMS pulse is determined by multiple factors whose combined contribution has yet to be fully quantified. Accounting for subject-specific pathway geometry allows TIDE to explicitly model and control a defined subset of the biophysical determinants of effective pathway engagement, reducing variability attributable to these factors and thereby providing a more controlled basis for investigating other sources of variability, such as the functional state of the brain at the time of stimulation.

Brain-state-dependent effects have been demonstrated in both motor and prefrontal cortex, where ongoing oscillatory activity can influence corticospinal excitability and TMS-induced plasticity (Gordon et al., 2021, 2022; Schaworonkow et al., 2019; Thies et al., 2018; Zrenner et al., 2018). Brain-state dependence therefore represents a genuine source of inter- and intra-individual variability.

However, the contribution of brain state may be difficult to isolate when stimulation dose is itself poorly controlled. Nominally identical stimulation may produce different levels of pathway engagement across participants, introducing variance that can obscure state-dependent effects. Tract-informed dosing and state-dependent stimulation therefore address complementary components of TMS variability: state-dependent approaches control when the pulse is delivered (Brancaccio & Miniussi, 2026), whereas TIDE improves control over the priming delivered to the relevant pathway. The observed response may thus reflect the interaction between the instantaneous susceptibility of the neural system and the effective pathway-level dose. Failing to account for the latter may lead apparent state-dependent effects to reflect, at least in part, biophysical differences in target engagement. This issue is particularly relevant in between-subject studies, but may also affect within-subject comparisons across targets, coil poses or sessions.

This reasoning is particularly relevant to TMS-EEG. Variability in TEPs cannot necessarily be attributed exclusively to fluctuations in cortical state or measurement noise, because nominally equivalent stimulation across participants may produce different effective doses and recruit the targeted neural elements or pathways to different extents (Dannhauer et al., 2024; Miniussi & Bortoletto, 2025; Numssen et al., 2024). By estimating stimulation intensity with respect to subject-specific tract geometry, TIDE may reduce this source of variability and improve the interpretability of electrophysiological and behavioural TMS outcomes.

The same principle may have clinical relevance. Clinical outcomes are likely to depend not only on the selected cortical coordinate and stimulation protocol, but also on whether the induced E-field effectively engages the pathway or network associated with the therapeutic target. In patients, this depends in the first instance on whether the pathway is still there to be engaged. Because TIDE operates on the individual tractogram rather than on a template, the estimate is conditioned on the residual architecture of that patient’s own bundle, and a pathway too disrupted or too sparsely reconstructed to support an estimate is flagged as such before stimulation is delivered. A tract-informed estimate could therefore complement anatomical and functional targeting by personalising the intensity required to engage the intended pathway. By providing an individualised estimate of pathway-level dose, TIDE could help reduce systematic under- or over-dosing across individuals and improve the consistency of target engagement in clinical settings as well.

### 4.1 Limitations

The present findings should be interpreted in light of several limitations concerning the behavioural validation, the assumptions of the biophysical model, and the tractography-based representation of pathway geometry.

First, the two validation arms are not equally informative about the shape of the dose-behaviour relationship. At P04, delivered doses occurred on both sides of the TIDE estimate, and the quadratic fit placed its maximum close to dose matching. This pattern is compatible with the proposed dose-matching relationship, but the small sample size precludes the definitive identification of a behavioural optimum. At P03, all participants were on the same side of the TIDE estimate, so the data cannot distinguish a two-sided dose-matching relationship from a monotonic association between bundle efficiency and behavioural responsiveness. The random-assignment analyses test the specificity of the subject-level TIDE mapping, not the behavioural effect of prospectively changing stimulation intensity. A within-subject design that samples multiple intensities around is therefore required to test the dose-response function directly.

Second, TIDE estimates depend on the accuracy of the underlying E-field model, including tissue segmentation, conductivity assumptions and coil modelling (Sections 2.5 and 2.11). Although errors shared by calibration and target simulations may partly cancel in the calibration ratio, region-specific modelling errors will not. TIDE estimates should therefore be interpreted as model-based quantities whose precision follows that of the underlying field simulation and is expected to improve with increasing head-model accuracy.

Third, the calibration inversion assumes that biological activation thresholds are approximately conserved between the calibration and target bundles (Section 2.9). However, axonal excitability depends on microstructural properties such as fibre calibre and myelination, which are not explicitly modelled by TIDE and cannot currently be recovered in vivo with the specificity required for bundle-wise calibration. Systematic differences in these properties between bundles could therefore introduce target-specific scaling errors in I_TIDE. Future advances in microstructural imaging may allow these factors to be incorporated more explicitly into the calibration.

Fourth, the framework assumes that the effect of stimulation on these pathways is mediated by direct activation of their white-matter axons, the D-wave-type mechanism in the corticospinal tract, rather than by indirect trans-synaptic recruitment; this direct-activation premise is the central physiological assumption on which the tract-level estimate rests. Even granting it, the activating function provides only a first-order approximation of how the induced field couples to an axon. As a linear, subthreshold quantity, it does not capture the full biophysics of axonal excitation, including membrane non-linearities, the dependence on pulse waveform and induced-current direction, temporal summation, and the enhanced excitability of terminals, bends and branch points (Aberra et al., 2020; Rattay, 1986, 1989). The resulting bundle-level estimate also depends on numerical and aggregation choices described in Sections 2.2–2.7. Although alternative aggregators are reported to assess sensitivity to these choices, they remain a source of model dependence.

Fifth, tractography streamlines are model-based reconstructions rather than direct representations of axons. Estimated bundle geometry depends on diffusion acquisition, segmentation, tracking and weighting procedures (Section 2.3; Supplementary Methods S1). Although the arc-length formulation reduces sensitivity to streamline sampling density (Section 2.4), other reconstruction-dependent sources of variability remain. Different tractography pipelines may therefore yield different TIDE estimates, and comparisons across studies should use consistent reconstruction procedures.

Finally, several practical constraints remain. RMT itself carries measurement variability, which propagates to the TIDE estimate through the calibration procedure (Section 2.9). The current framework addresses single-pulse dose (i.e., intensity) and does not model the protocol-dependence, or temporal accumulation dose relevant to repetitive stimulation. In addition, some recommended intensities may fall outside the technical and safety range deliverable by a given stimulator (Rossi et al., 2021). Finally, the present behavioural validation is limited to 19 participants, one task and two endpoints of a single tract, and its generalisability across pathways, paradigms and cohorts remains to be established.

## 5. Conclusions

TIDE provides a tract-informed framework for translating motor-threshold calibration into individualised stimulation intensities for non-motor white-matter targets. By combining an observable physiological anchor with subject-specific E-field modelling and tractography, the method moves beyond conventional dosing based on cortical E-field magnitude alone and accounts for how the induced field interacts with the architecture of the pathway intended to be engaged.

In an independent behavioural cohort, smaller deviations between the delivered intensity and the TIDE recommendation were associated with larger site-specific behavioural effects, providing retrospective evidence that TIDE-derived dose differences are functionally relevant.

Furthermore, across thirteen white-matter targets, the substantial inter-individual variability in the TIDE-recommended intensity indicates that pathway-specific dose can rarely be reduced to a single group-level value. Because the calibration anchor was held constant across participants in that analysis, this variability is geometric in origin, and it challenges the assumption that a fixed percentage of RMT delivers a comparable effective dose across cortical regions.

Together, these results support TIDE as a practical framework for investigating and reducing variability in effective pathway engagement across TMS applications by providing an anatomically grounded estimate of stimulation dose. The approach may strengthen experimental control in cognitive and TMS-EEG studies and complement anatomical and functional targeting in clinical protocols.

## 7. Intended use and regulatory status

TIDE is intended for research use only. It is not a medical device and must not be used to diagnose or treat patients or to guide clinical decisions. The software has not been evaluated by any regulatory authority and holds no regulatory clearance: neither CE marking under the European Medical Device Regulation nor approval from the United States Food and Drug Administration. Any clinical use would require independent validation and the appropriate regulatory authorisation. The same Research-Use-Only statement is provided in the --version and --help CLI option flags, and in the software’s documentation.

## 8. Data Availability

TIDE is open-source software distributed under the GNU General Public License v3.0 or later (GPL-3.0-or-later). The source code is hosted on GitHub (https://github.com/marcotag93/TIDE), and the Python package is distributed through the Python Package Index (PyPI) (https://pypi.org/project/tide-pipeline/). The software version associated with this manuscript is archived on Zenodo under DOI <u>10.5281/zenodo.22019737</u>. The analysis code, pre-registration record, derived data, and other materials required to reproduce the analyses reported in this manuscript are archived separately on Zenodo (10.5281/zenodo.22019794).

