## Supplementary Methods and Figures for "TIDE: Tractography-Informed Dose Estimation for individualised TMS intensity"

#### S1 — Tractogram requirements and recommended preprocessing

For the diffusion acquisition, a multi-shell protocol with at least two non-zero b-values (typically b = 1000 and b = 2000–3000 s/mm², with 60 or more directions on the outer shell) gives the angular resolution required to resolve crossing fibres in the lateral association tracts and the corona radiata. Standard preprocessing (denoising, Gibbs unringing, eddy-current and motion correction, bias-field correction, registration to the T1-weighted reference) is performed in subject space, so the streamlines remain coregistered to the mesh that drives the FEM solve. Fibre orientation distributions are estimated with constrained spherical deconvolution, in its multi-shell multi-tissue form when more than one shell is available, at maximum harmonic order $l\_max$= 6–8.

Bundle reconstruction is performed with TractSeg (Wasserthal et al., 2018), using its default U-Net segmentations of the bundle mask, the start and end regions, and the tract orientation maps. For each bundle, probabilistic tracking is run inside the bundle mask with the TOM as a directional prior, with a minimum streamline length of 50 mm, a maximum length appropriate to the bundle (typically 200–250 mm for long association tracts), a step size of 0.5–1 mm, and a target of at least 5000 streamlines per bundle before any filtering. The step-size recommendation matters: the AF along a streamline is the along-fibre spatial derivative of the parallel field component, and a step size larger than 2 mm under-samples that derivative in Eq. (1) even after the arc-length resampling and physical Gaussian smoothing of Section 2.4.

A manual refinement pass sits between tracking and SIFT2. Probabilistic tracking inside a TractSeg mask recovers the bundle at the cost of false-positive streamlines: fibres that leave the tract at a crossing region, loop back on themselves, or terminate in a neighbouring gyrus. These are geometrically well-behaved curves, so they pass the four-point cutoff and the angular-deviation filter of Section 2.4 and enter Eq. (3) with the same status as the rest of the bundle. Where their course is misaligned with the induced field they depress the per-streamline threshold distribution; where it happens to be favourably aligned they inflate the upper tail on which the aggregator of Eq. (4) is computed. We therefore refined every bundle by virtual dissection in TractEdit (v3.4.7; (Tagliaferri & Cattaneo, 2026), with one operator removing streamlines whose course was inconsistent with the accepted anatomy of the tract, and the same criteria applied to the target bundles and to the calibration corticospinal tract. The cortical target coordinates that seed coil optimisation and grid search were read from the refined bundles in the same session and in the same RASMM frame as the head model, so the coordinate that defines the ROI of Section 2.2 and the streamlines that populate it come from a single tractogram. Refinement was completed before any E-field simulation, so the edits could not be informed by the resulting dose estimates, and before SIFT2, so the per-streamline weights described next were estimated on the refined tractogram and correspond one-to-one to the streamlines that TIDE loads.

SIFT2 (Smith et al., 2015) is then run on the per-bundle tractogram and the resulting per-streamline weights are passed to TIDE through subject.files.weights_cst for the calibration bundle and subject.files.weights_target for the target bundle; the weighted aggregator of Section 2.7 picks them up automatically.

Two requirements are bundle-specific. Cortical termination is essential, because the contiguous-activation criterion of Section 2.6 evaluates the AF on the streamline segment that lies inside the ROI sphere centred on the cortical target. TractSeg's ending-region segmentation should be used to constrain the streamline endpoints to the cortical or immediately subcortical white matter; bundles whose terminations are restricted to deep white matter will report low AF magnitudes that reflect the cortical-reach limit of the coil, not a property of the bundle itself. Streamline density inside the ROI should be at least 500 streamlines, so that the empirical 95th percentile of the per-streamline threshold distribution is stable; bundles with sparse coverage of the ROI should be either reseeded with a higher target count or excluded from analysis. The pipeline logs the analysed streamline count for every bundle, so a degradation in coverage is visible in the standard output.

#### S2 — Field-solver threading and field-sample matching

Worker processes that host the FEM solver are configured single-threaded: `OMP_NUM_THREADS`, `MKL_NUM_THREADS`, `OPENBLAS_NUM_THREADS`, and `NUMEXPR_NUM_THREADS` are all set to 1 in the parent before the worker pool spawns, so that each child inherits single-threaded BLAS at interpreter start-up, before it imports NumPy or SimNIBS, and the cumulative thread count is bounded by the number of Python workers rather than by the product with the OpenMP pool.

The values returned by the sampler are matched back to the requested streamline coordinates with a 0.1 mm KD-tree tolerance before use. Coordinates returned in their original order take a direct path; a reordering or a small perturbation of the returned points is realigned against the request, and a non-bijective match, in which two requested points share one nearest neighbour, raises rather than proceeding on corrupted output. A streamline point with no returned value is filled with `NaN` rather than zero, so that a missing sample is detectable instead of silently deflating the along-fibre gradient. The unmatched count is logged, and a loss above 1 % of requested points raises, on the assumption that it signals a genuine coordinate or ordering mismatch. Streamlines carrying a non-finite AF inside the ROI are dropped before aggregation (Section 2.7), so that a sub-1 % miss degrades the bundle metric gracefully rather than poisoning it.

#### S3 — Weighting and surface-constraint modes

The cross-streamline aggregator of Section 2.7 admits two orthogonal extensions that are exposed as configuration options of the pipeline: a weighting scheme that compensates for streamline density bias, and a surface filter that restricts the aggregation to a band of tissue around the grey-white interface. The two extensions are configured independently. The surface filter acts once per run, upstream of the per-streamline threshold scan, so a given run is either surface-constrained or it is not; changing that setting requires a second run. Within a run, the weighted and unweighted aggregates are computed together from the same array of per-streamline thresholds and reported side by side.

The unweighted mode is the empirical estimator of Eq. (4) and is the safe default for tractograms without weight files. The SIFT2-weighted mode (Smith et al., 2015) replaces the empirical percentile and median by their weight-aware counterparts, with each streamline contributing a fraction of the cumulative weight rather than a single count; weights are accepted either as a one-per-line text file (the SIFT2 standard output) or as a NIfTI scalar volume sampled along each streamline as the mean voxel value. A supplied weight file is validated on load: a weight set that fails to read, or that carries non-finite, negative, or zero-total-mass values, aborts the run with an explicit error rather than silently reverting to uniform weighting, as does a weight vector shorter than the largest streamline index the analysed bundle references. Uniform weighting is used only when no weight file is configured. Weights are indexed by original streamline id, so the weight file must be estimated on the same tractogram that TIDE loads; a weight vector left over from an earlier, larger version of the bundle is longer than the file it is paired with, is not detected by the load-time checks, and would give each streamline another streamline's weight. Because streamlines can be dropped downstream of that point (short stubs, Frenet failures, or the angular filter of Section 2.4), the original streamline indices are carried through the drop chain so that the weight vector is realigned to the surviving set before aggregation, and each retained streamline keeps its own SIFT2 weight rather than inheriting a neighbour's.

The surface-constrained mode uses a FreeSurfer white-matter surface (`lh.white` or `rh.white`; Fischl, 2012) loaded into a KD-tree; streamline points further than `options.gwi_threshold_mm` from the surface (3 mm default) are excluded from the per-streamline threshold computation. The mode is enabled by supplying `subject.files.surface`; when no surface is configured the cutoff is unused. Enabling the surface filter alongside SIFT2 weighting is appropriate when bundles of markedly different anatomical depth are compared. This is the concurrent setting of two independent switches, not a distinct analysis mode with its own code path.

Every run reports the weighted and the unweighted aggregate side by side, each carrying a label that records the provenance of the weights actually used — the configured weight file, or `Uniform` when none is supplied. The surface constraint is a property of the run rather than of a row, and is recorded in the YAML configuration snapshot written beside the report. Downstream consumers default to the weighted variant when SIFT2 weights are supplied and to the unweighted variant otherwise, so a tractogram without weights yields an unweighted estimate and one with weights yields a weighted estimate without manual configuration changes. A degradation in the weighted variant relative to the unweighted variant on the same tractogram is diagnostic of a mismatch between the SIFT2 weights and the analysed streamline subset.

#### S4 — Output schema and artefacts

Output directory layout, file naming, and the column order in `TIDE_Results_<target>.txt` and `TIDE_grid_results.csv` are preserved across patch and minor releases. Each text report is accompanied by a same-stem JSON sidecar, which preserves the exact report lines together with parsed sections and any structured metadata, and a self-contained HTML rendering; both are reporting-only and add no quantity not already present in the text report. Breaking changes are recorded by a major-version bump of `tide.__version__`, which is written into the header of every report so that any reported number is traceable to the producing code version. Downstream tools that parse the reports by name and column position therefore remain functional across releases.

Worker assignment follows the structural symmetry of the pipeline. The M1/CST branch — coil optimisation, FEM simulation, streamline sampling, and AF computation — runs in one process; the equivalent target-side branch runs concurrently in a second. Process isolation uses Python's `multiprocessing.spawn` context; the single-threaded BLAS configuration that bounds the worker thread count is described in Supplementary Methods S2. A deliberate read of the head mesh on the main process before workers spawn warms the operating-system disk cache, after which the workers load the same file at near-RAM speed. The two branches join at a synchronisation barrier immediately before the calibration ratio is evaluated.

The run writes to <derivatives>/sub-<id>/TIDE_<target_label>/. Its primary artefact, TIDE_Results_<target>.txt, is generated from the shared template in core/io.build_estimation_summary_lines and contains, in this order, a run header, the configuration block, the M1 calibration block (measured RMT, coil matrix, pose-quality control, weighted and unweighted CST efficiency, RMT dI/dt, and biological threshold), the target estimation block (target coordinates, scalp position, coil matrix, pose-quality control, and weighted and unweighted target efficiency), the geometric-analysis block, the self-validation metrics, and the primary results table containing the raw and clamped intensity estimates, clamp flags, SEI, and calibration multiplier. An additive Aggregator Sensitivity block follows the primary results when sensitivity data are available; these diagnostic values do not alter the dose reported in the primary results table. The companion artefacts are the AF-annotated tractograms in TRK format with signed AF and segment-length scalars attached at each streamline point, the optimisation matrices in human-readable form, the SimNIBS simulation directories, and a snapshot of the YAML configuration used by the run. The signed activating function is retained in the streamline (TRK) scalars, whereas the volumetric NIfTI overlays store |AF|, because most viewers (FSLeyes, MRIcroGL, ITK-SNAP) render scalar volumes without a diverging colour map by default; keeping the signed representation on the streamlines and the magnitude representation in the volumes preserves both downstream uses.

A self-validation block runs at no additional FEM cost. The already-computed M1 FEM E-field is sampled along the target tractogram; the contiguous-threshold scan of Section 2.6 is re-applied; and the block reports AF_target at the M1 pose, the optimisation gain over that pose, and the I_TIDE that the M1 pose would prescribe for the target bundle. This tests internal consistency between the calibration and target arms without invoking a second FEM solve. The scan used here is a simplified variant of Eq. (3): it evaluates the criterion on the in-ROI midpoint magnitudes against the native segment lengths, without the adjacent-midpoint averaging or the minimum-coverage gates of Section 2.6. The cross-streamline aggregation is identical. The block is therefore a consistency diagnostic and is deliberately not reported as an alternative estimate of $AF\_target$; the values that enter the calibration inversion of Eq. (5) come exclusively from the Section 2.6 estimator.

The report carries two quality-control diagnostics. The first is a coil-pose plausibility check that compares the final `matsimnibs` axis against the local outward scalp normal and flags inferior or upward-firing skull-base poses. The second is an alignment diagnostic that compares midpoint streamline tangents with midpoint E-field vectors. Neither diagnostic modifies the optimisation, the AF, the SEI, or the computed intensity. The alignment diagnostic is purely advisory. The plausibility check is advisory for a user-supplied 4×4 matrix, which is treated as a specialist override and only warned, but it acts as a dose-eligibility gate for an automatically optimised pose: a pose that fails the check stops the estimation run, and a failing grid point is recorded as a failure and excluded from the dose and ranking summaries, so that a pathological automatic pose never yields a trusted dose.

Each candidate is processed by a worker pool whose size is selected dynamically from the available CPU and memory budget. Worker identifiers are assigned through file-locked claim files in the output directory, so that a crashed worker can be replaced without renumbering the remaining tasks.

Run-wide artefacts include TIDE_grid_results.csv, with one row per successful grid point. Its core columns record the per-point cortical and scalp coordinates, coil matrix, raw and clamped weighted and unweighted intensities, clamp flags, SEI, within-run SEI percentile rank, and calibration multipliers; additive columns report the weighted and unweighted raw intensity obtained with each aggregator in the sensitivity analysis;`TIDE_Grid_Summary_<target>.txt`, a per-target overview listing the best and worst points by each reported quantity together with the distribution of clamp flags; `grid_points_mask.nii.gz`, a binary mask of candidate positions in T1w space; and a configuration snapshot.

Per-point artefacts under `simulations/grid_P<N>/` mirror the estimation output. Each candidate carries a `TIDE_Results_<target>.txt` generated from the same template as the standalone report and is therefore bit-equivalent in content to a single-estimation run. Alongside the report, the candidate's coordinates, scalp coordinates, and 4×4 `matsimnibs` matrix are written into a reproducible YAML configuration with the target section pre-filled, so that the estimation workflow can be re-run on any candidate without manual editing. A visualization/ subdirectory holds the cortical-surface deliverables. Three scalar NIfTI volumes are written in T1w space: grid_mso_map.nii.gz carries I_TIDE,clamped at each grid point, grid_mso_raw_map.nii.gz carries the unbounded I_TIDE of Eq. (5), and grid_mso_flag_map.nii.gz carries the clamp regime as an integer code (1 = WITHIN_RANGE, 2 = CLAMPED_LOW, 3 = CLAMPED_HIGH, 4 = DEVICE_LIMITED, 0 = background). The two intensity volumes coincide wherever the estimate falls inside the clamp envelope and diverge only where the flag departs from WITHIN_RANGE, so neither quantity overwrites the other. A label sidecar (grid_mso_labels.tsv) lists both intensities and both clamp flags per candidate for FSLeyes and MRIcroGL; its first eight columns are held fixed for backward compatibility and the remaining fields are appended. Candidates whose inversion failed carry no row in TIDE_grid_results.csv and appear in none of the three volumes. A self-contained interactive HTML viewer (grid_interactive.html) renders the cortical surface and the candidate positions in a standard browser without external dependencies; it colours the candidates by either quantity under a labelled colour bar, with the raw estimate shown by default.

#### S5 — Reference implementation, distribution, and testing

The TIDE package imports from the SimNIBS 4.5+ Python environment that also hosts the FEM solver of Section 2.5. Three-dimensional rendering is optional and is provided by an extra (`pip install tide-pipeline[viz]`) that adds PyVista and VTK; a run without it produces every numerical and tabular output and omits only the rendered figures. Every direct runtime dependency and optional dependency is pinned to an exact version in the PEP 621 project metadata, and the build backend is fixed to hatchling 1.27.0. The repository additionally carries a uv lockfile that freezes the transitive dependency graph used for the locked development and release environment. The lockfile is not consumed by a standard PyPI/pip installation; reproducibility of the numerical runtime is therefore additionally protected by the SimNIBS-environment version checks described below.

The package is distributed on PyPI as *tide-pipeline*, while the Python import package and command-line entry point are both named *tide*. SimNIBS is a separately installed system dependency and is never resolved from PyPI by TIDE. The recommended PyPI installation uses an isolated launcher environment, for example *pipx install --python 3.11 tide-pipeline* or *uv tool install --python 3.11 tide-pipeline*, followed by *tide --bootstrap*. The bootstrap command locates the SimNIBS Python explicitly, verifies its numerics-critical dependency versions against TIDE's pinned versions, and installs the matching TIDE release into that interpreter. A conventional python -m pip install tide-pipeline remains supported when installation into the current interpreter is intentional. From a source checkout, the recommended development installation is python install.py --simnibs-env --editable. The installed package also provides *tide --init-config config.yml*, which materialises the canonical annotated configuration template from package data without requiring SimNIBS to be initialised.

Platform support follows SimNIBS (Linux, macOS, and Windows on 64-bit architectures).

The user-facing entry point is the \`tide\` console script, registered in \`pyproject.toml\` and dispatched through \`tide.cli.main\`. The dispatcher accepts a \`--config\` flag pointing to a single YAML file that drives all workflows and a \`--verbosity\` {quiet, standard, verbose} flag for log-level control. Each tagged release carries a semantic version defined in src/tide/__init__.py, exposed at runtime as tide.__version__ and propagated into the package metadata by hatchling. The release workflow verifies that this version agrees with CITATION.cff and, for a production release, with the GitHub release tag. The software version is also displayed by tide --version and in the HTML report sidecar. Backward-incompatible changes to the output schema, the YAML schema, or the calibration arithmetic of Eq. (5) trigger a major-version bump.

When the final coil pose is a full 4×4 \`matsimnibs\` matrix, whether supplied in the configuration or returned by optimisation, the SimNIBS field result is stored in a content-addressed cache keyed on the head mesh, coil model, matrix, dI/dt, requested fields, and runtime versions. A later matching run restores the mesh and its companions byte-for-byte and continues through the unchanged sampling and AF path, which cuts the FEM step from minutes to seconds without altering any downstream value; a corrupt or mismatched entry is treated as an ordinary cache miss and recomputed. The cache is out-of-band and regenerable, holds no analysis-grade data, and can be relocated or disabled from the configuration or the command line.

The test suite under tests/ provides regression coverage for the activating-function computation (Section 2.4), the contiguous-activation threshold (Section 2.6), the cross-streamline aggregator and its sensitivity diagnostics (Section 2.7), the clamp arithmetic (Section 2.9), configuration initialisation and environment bootstrapping, and the separation of raw, clamped, and clamp-regime maps in the grid visualisation (Supplementary Methods S4). The SimNIBS-independent suite is executed by continuous integration on Python 3.11 and 3.12 for pushes and pull requests. The release workflow builds the source and wheel distributions, validates the package metadata and README rendering, installs the built wheel in a clean environment, and smoke-tests the installed package, console entry point, and bundled configuration template. A manual run from an exact version tag publishes only to TestPyPI. Publishing a GitHub Release from the same tag rebuilds the distributions, verifies that they are byte-identical to the TestPyPI artifacts, and then publishes them to PyPI. The first is the M1 identity case, in which the calibration and target bundles are the same bundle at the same pose and the inversion of Eq. (5) is required to return the measured RMT. The second is a set of analytic checks on the activating function itself: a synthetic field with a known along-fibre gradient is substituted for the FEM output on straight and curved fibres, and the recovered AF is required to match the analytic value in SI units at both boundaries and to remain invariant under resampling of the fibre between 0.25 and 2.0 mm, under reversal of streamline direction, and under up- and down-sampling of a curved fibre. Both are pure-Python and require no SimNIBS installation.

#### S6 — Conductivity model: anisotropy bias mechanism

The bias structure is straightforward. Anisotropic conductivity tensors are conventionally constructed by linear scaling, volume normalisation, or volume-anisotropy normalisation of the DTI tensor field (Güllmar et al., 2010; Tuch et al., 2001). Each of these mappings is a deterministic function of the same eigenstructure that the streamline-tracking algorithm uses to grow streamlines. The resulting conductivity tensor therefore correlates with the local fibre orientation by construction: regions of high parallel conductivity coincide with regions where the streamlines align with the field gradient, and regions of low transverse conductivity coincide with the bends in the streamlines. The activating function on a streamline depends, by Eq. (1), on the along-fibre gradient of the dot product of the field with the streamline tangent; this dependency is amplified by an anisotropic conductivity that is itself aligned with the streamline. The amplification is not the same on the calibration bundle (CST) and on an arbitrary target bundle, because the two bundles have different orientation distributions and different volumetric overlap with the high-conductivity regions of the head. The ratio AF_CST/AF_target inherits this differential amplification, and the inferred I_TIDE of Eq. (5) becomes a function of the DTI-to-conductivity mapping in addition to the bundle anatomy.

The published literature on TMS E-field modelling supports the design choice on independent grounds. The cortical E-field at the gyral surface is modestly sensitive to white-matter anisotropy under DTI-derived conductivity tensors. Opitz et al., 2011 reported peak-to-peak changes of approximately 5–10 % in the surface field magnitude when comparing isotropic and anisotropic head models on the same subject; Thielscher et al., 2011 reached similar conclusions on a different mesh. Saturnino et al., 2019 confirmed that the field magnitude near the cortical hotspot is robust to the conductivity model and noted that the dominant residual uncertainty is the DTI-to-conductivity mapping itself, for which no consensus exists. The order of magnitude of the anisotropic correction at the cortical surface is therefore smaller than the order of magnitude of the across-bundle differences that the TIDE calibration is designed to capture, and the correction is paid for by a bias that does not cancel in the ratio.

#### S7 — Coil-pose specification and field sampling

The reference finite-element solve of Section 2.5 fixes the coil pose and then samples the induced field along each streamline; the pose-specification routes and the sampler are described here.

The coil pose is specified by one of two routes. The first route accepts a user-supplied 4×4 `matsimnibs` matrix that encodes coil position and rotation in mesh coordinates; the matrix is applied without further optimisation, which is the appropriate behaviour when calibration or target must reproduce a fixed pose, for example a coil position recorded in a previous experimental session by neuronavigation or by a previous TIDE-based or SimNIBS-based simulation. The second route accepts a cortical target coordinate and runs the auxiliary dipole method optimisation (ADM, Gomez et al., 2021), which evaluates a grid of candidate scalp positions and handle rotations on the user-supplied target. The direct method (`opt_struct.TMSoptimize` with `method='direct'`) is available as a user-selectable alternative to ADM, enabled by setting `options.adm_optimization: false` in the configuration; the pipeline does not switch between the two methods automatically. The coil-to-scalp distance is fixed at 4 mm by default, which matches the padding typically present between coil casing and scalp surface during routine TMS sessions; the value is configurable through the `coil.coil_distance_mm` field of the input configuration.

When the user supplies only a cortical target, the scalp entry point is computed by ray–triangle intersection (Möller & Trumbore, 1997) between a ray from the brain centroid through the cortical target and the scalp surface mesh. When no handle reference is supplied, the default orientation projects the anterior direction onto the scalp tangent plane and rotates it 45° toward the medial direction, with the medial sense inferred from the sign of the scalp position's lateral coordinate; this reproduces the conventional handle orientation for left-hemisphere motor-cortex stimulation and extends to the right hemisphere without manual intervention.

The E-field is sampled at every streamline point by tetrahedral interpolation on the FEM solution. Streamlines are kept in RASMM space (Section 2.2) and the SimNIBS mesh is loaded in the same space, so no resampling of streamline coordinates is required before interpolation. The sampling routine, `interfaces.sampling.sample_field_at_coordinates`, returns a list of arrays, one per streamline, each of shape (Npoints, 3) with field components expressed in V/m per A/μs (intrinsic efficiency form, given the unit reference dI/dt). These field arrays are then consumed by the AF computation of Section 2.4.

#### S8 — Cross-streamline aggregator: robustness properties and alternatives

The bundle-level aggregator adopted in Section 2.7 is the median of the top five per cent of the per-streamline thresholds T_i (Eq. 4). Its robustness properties, and the alternative aggregators evaluated on the same per-streamline threshold distribution and reported alongside it, are set out here.

The median-of-top-five-per-cent aggregator has three properties that motivate its use as the bundle-level statistic. First, the aggregate is concentrated on the streamlines that carry the physiologically informative signal, that is, streamlines for which a contiguous supra-threshold activation length actually exists; streamlines below the 95th percentile of the per-streamline threshold distribution, including stubs that pass the four-point cutoff but contribute negligible activation, do not enter the aggregate. Second, the median operation inside the retained subset stabilises the estimator against the upper tail of the per-streamline distribution, which is the region most affected by residual tractography noise and by streamlines that graze the ROI without supporting a coherent activation length. Third, the percentile-then-median construction reduces sensitivity to streamline density: doubling the number of streamlines in a tractogram of fixed anatomical content leaves Q0.95, and the median of the retained subset approximately invariant, since both are quantile statistics of the same underlying distribution.

The three quantile properties set out above motivate this default analytically, but the choice remains a modelling decision rather than a measured optimum, and the pipeline does not hedge it: the median of the top five per cent is the statistic that drives the calibration inversion, while five alternative aggregates — the median of the top one per cent, the 95th and 90th percentiles, the median, and the mean, each in weighted and unweighted form — are computed on the same per-streamline threshold distribution and written to the estimation and per-grid-point reports as a sensitivity table; the grid-results CSV additionally carries the corresponding weighted and unweighted raw-intensity values in additive columns (Supplementary Methods S4). The sensitivity of $AF\_CST/AF\_target$ to the aggregation rule is nonetheless recoverable without re-running the field solve, because the signed activating function and the native segment lengths are retained per streamline in the released TRK outputs (Supplementary Methods S4). Applying Eq. (3) to those scalars reproduces the per-streamline threshold distribution $T\_i$ for each bundle, on which any alternative aggregator can be evaluated offline. We regard these alternatives as an in-pipeline sensitivity diagnostic rather than competing estimates, and we make no claim that the reported ratio is invariant to the choice.

#### S9 — Intensity clamp and QC-flag arithmetic

Eq. (5) is linear in RMT, so an extreme value of the AF ratio can produce an I_TIDE outside the configured reporting interval or beyond the maximum output of the stimulator. TIDE therefore retains the raw inversion and additionally reports an operationally bounded value. The upper bound is I_ceil = min(r_c · RMT, 100%), where 100% is the physical maximum stimulator output, and the lower bound is I_floor = min(r_f · RMT, I_ceil), so the configured floor can never exceed the programmable ceiling. The defaults r_f = 0.70 and r_c = 1.40 are configurable through options.mso_floor_ratio and options.mso_ceiling_ratio, and the lower bound is disabled by setting r_f = 0. These bounds are reporting and operational constraints rather than physiological safety thresholds; they do not change the raw model estimate or any upstream activating-function quantity.

Both I_TIDE and I_TIDE, clamped are reported in every output, together with a flag value that records the regime of Eq. (S1). WITHIN_RANGE, CLAMPED_LOW, CLAMPED_HIGH, and DEVICE_LIMITED partition the finite case according to which leg of Eq. (S1) is active and, at the ceiling, which of the two bounds binds. ESTIMATION_FAILED is emitted instead when the raw inversion is not finite, which happens when AF_target is zero or undefined; in that case both intensities are written as missing values and the row is excluded from grid aggregates and rankings, so that a failed inversion is never carried forward as a clamped dose. The clamp never propagates to the multiplier k, the SEI, or any upstream quantity.

From Eq. (7) and the floor leg of Eq. (S1), the closed-form relation between the CLAMPED_LOW regime and the SEI is

$$\text{CLAMPED\_LOW} \Leftrightarrow \text{SEI} > 1 / rf$$

At the default mso_floor_ratio = 0.70, the lower clamp engages for SEI > 1.43; the symmetric upper-clamp condition engages for SEI < 0.71 at the default ceiling of 1.40.

#### S10 — Random-assignment tests of subject-specific TIDE estimates

We used two permutation analyses to test whether the observed negative correlation between TIDE–delivered dose mismatch and behavioural response depended on the correct subject-specific pairing of TIDE estimates and participants, or could arise when the observed TIDE estimates were randomly reassigned among participants. These analyses were secondary robustness tests and did not replace or modify the pre-specified behavioural analysis described in Section 2.12.

P03 and P04 were analysed separately, and permutations were generated independently for the two sites. The observed sham-corrected behavioural shift and the experimentally delivered intensity were held fixed for every participant throughout.

In the absolute-intensity permutation analysis, the raw I_TIDE values expressed in %MSO were randomly reassigned among the 19 participants within each site. For each permutation, the relative mismatch was recalculated as

$$m_i^{perm} = \frac{\left|I_{delivered,i} - I_{TIDE,i}^{perm}\right|}{I_{TIDE,i}^{perm}}$$

and Spearman's rank correlation between $m^{perm}$ and the observed $\Delta RI$ was computed. This procedure preserves the empirical distribution of TIDE-recommended absolute intensities but breaks their subject-specific assignment. Because an absolute TIDE intensity is expressed on an individual RMT-derived stimulation scale, this null model also disrupts the natural coupling between the TIDE estimate and that participant's absolute stimulation scale.

The second analysis therefore randomised SEI directly. In the validation cohort,

$SEI = \frac{AF_{target}}{AF_{CST}} = \frac{I_{delivered}}{I_{TIDE}}$Because I_delivered is the RMT × 120% calibration anchor used for the retrospective inversion. Consequently,

$$\frac{\left|I_{delivered} - I_{TIDE}\right|}{I_{TIDE}} = |SEI - 1|$$

Randomising SEI therefore removes the participant-specific absolute RMT/%MSO scale from the randomised predictor and tests the specificity of the target-to-CST efficiency assigned by TIDE. For each permutation, the 19 observed SEI values were reassigned among participants within the site, $m^{perm} = |SEI^{perm} - 1|$ was calculated, and its Spearman correlation with the fixed behavioural outcome was recorded.

Each null distribution comprised 100,000 random assignments. The directional alternative was the same as in the primary analysis: stronger TIDE-behaviour relation corresponds to a more negative Spearman coefficient. Directional permutation probabilities were therefore calculated as

$$p = \frac{1 + \sum_{b=1}^{B} \mathbf{1}\left(\rho_b \leq \rho_{obs}\right)}{B + 1}$$

With $B = 100{,}000$. The correction prevents zero-valued Monte Carlo permutation probabilities. Fixed random seeds were used for reproducibility. Behavioural values were never permuted. The two analyses define different null hypotheses and their permutation probabilities are therefore not expected to be identical.

### Supplementary Figures

#### S1 — Absolute I_TIDE %MSO permutations

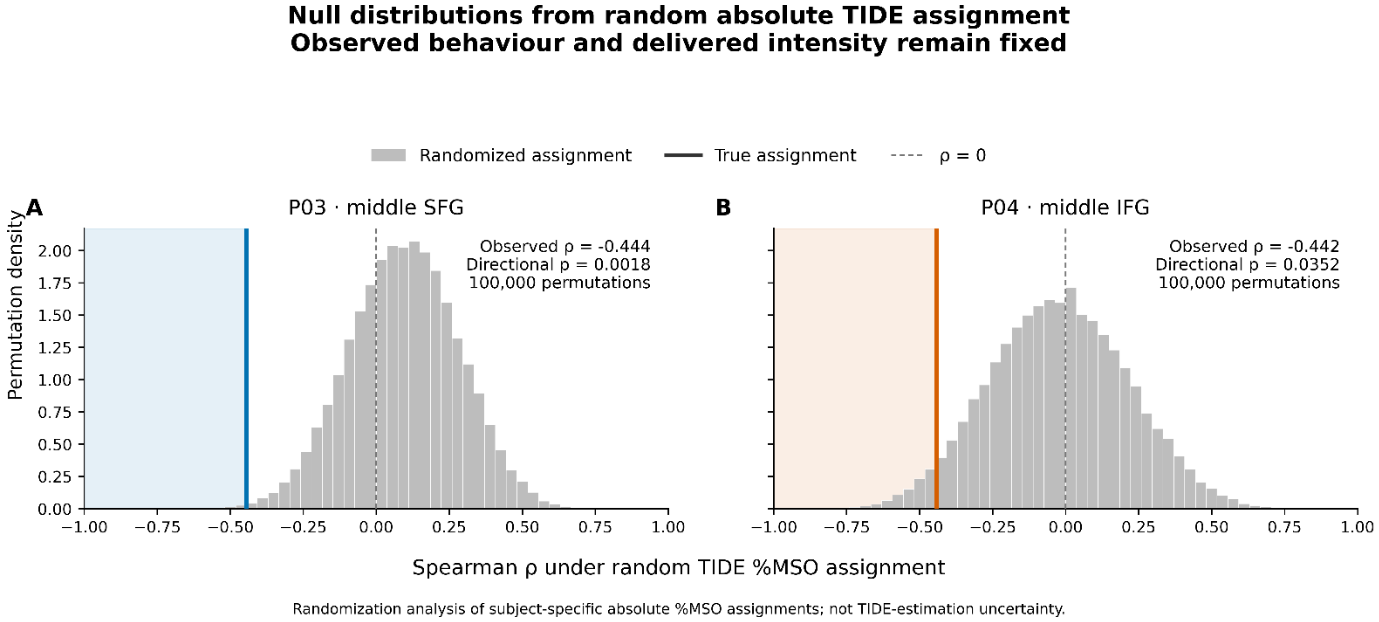


**Supplementary Figure S1. Random reassignment of subject-specific absolute TIDE intensities.** Null distributions were generated by randomly reassigning the observed raw TIDE-recommended intensities in %MSO among participants while keeping each participant's delivered intensity and sham-corrected behavioural response fixed. For each random assignment, the relative mismatch between delivered and reassigned TIDE intensity was recalculated and correlated with the observed behavioural shift using Spearman's . Grey histograms show the null distributions from 100,000 independent permutations within each site; coloured vertical lines show the correlations obtained with the true subject-specific TIDE assignments, and shaded areas mark the directional tails at least as negative as the observed coefficients. **(A)** P03, middle SFG: observed $\rho = -0.444$, directional permutation $p = 0.0018$. **(B)** P04, middle IFG: observed $\rho = -0.442$, directional permutation $p = 0.0352$. This null model randomises the absolute TIDE recommendation and therefore also disrupts its coupling to the participant-specific absolute stimulation scale. SFG, superior frontal gyrus; IFG, inferior frontal gyrus; MSO, maximum stimulator output.

### S2 — SEI permutations

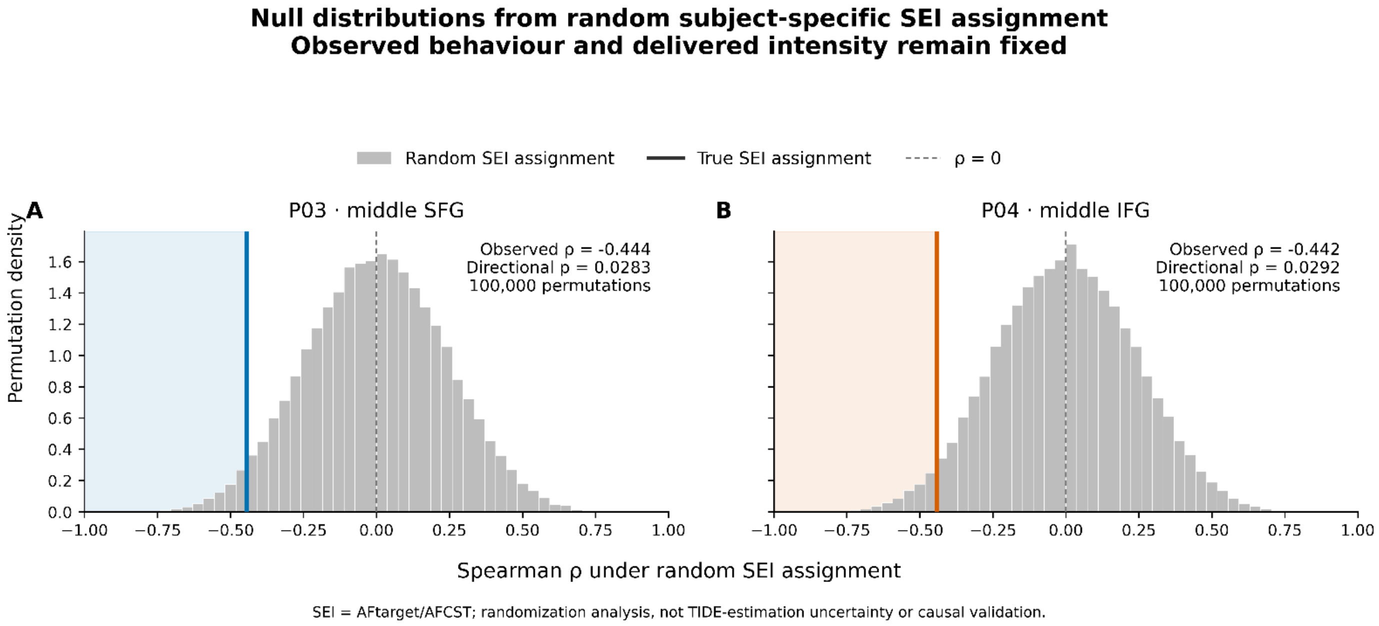


**Supplementary Figure S2. Random reassignment of subject-specific Stimulation Efficiency Index values.** SEI values were randomly reassigned among participants while the observed behavioural responses and delivered intensities remained fixed. For each permutation, the mismatch was calculated directly as $|SEI - 1|$ and correlated with the observed sham-corrected behavioural shift. Because the absolute stimulation anchor cancels from this quantity, this analysis randomises the subject-specific target-to-CST efficiency without randomising the participant's absolute RMT/%MSO scale. Grey histograms show the null distributions from 100,000 independent permutations within each site; coloured vertical lines show the correlations obtained with the true subject-specific SEI assignments, and shaded areas mark the directional tails at least as negative as the observed coefficients. **(A)** P03, middle SFG: observed $\rho = -0.444$, directional permutation $p = 0.0283$. **(B)** P04, middle IFG: observed $\rho = -0.442$, directional permutation $p = 0.0292$. SFG, superior frontal gyrus; IFG, inferior frontal gyrus; SEI, Stimulation Efficiency Index.
